# Sequential antibiotic therapy in the lab and in the patient

**DOI:** 10.1101/2022.06.17.496526

**Authors:** Christin Nyhoegen, Hildegard Uecker

## Abstract

Laboratory experiments suggest that rapid cycling of antibiotics during the course of treatment could successfully counter resistance evolution. Drugs involving collateral sensitivity could be particularly suitable for such therapies. However, the environmental conditions *in-vivo* differ from those *in-vitro*. One key difference is that drugs can be switched abruptly in the lab, while in the patient, pharmacokinetic processes lead to changing antibiotic concentrations including periods of dose overlaps from consecutive administrations. During such overlap phases, drug-drug interactions may affect the evolutionary dynamics. To address the gap between the lab and potential clinical applications, we set up two models for comparison - a ‘lab model’ and a pharmacokinetic-pharmacodynamic ‘patient model’. The analysis shows that in the lab, the most rapid cycling suppresses the bacterial population always at least as well as other regimens. For patient treatment, however, a little slower cycling can sometimes be preferable if the pharmacodynamic curve is steep or if drugs interact antagonistically. When resistance is absent prior to treatment, collateral sensitivity brings no substantial benefit unless the cell division rate is low and drug cycling slow. By contrast, drug-drug interactions strongly influence the treatment efficiency of rapid regimens, demonstrating their importance for the optimal choice of drug pairs.

## 1 Introduction

Resistance of bacteria to antibiotic treatment is a tremendous problem for health care world-wide, caused by the fact that bacteria, as all organisms, evolve and adapt to their environments. Such an evolution-caused problem calls for a solution based on evolutionary principles. Evolution-informed medicine aims to develop sustainable treatment strategies that prevent the selection of resistance during treatment [1–3]. One such strategy is sequential therapy, a multi-drug regimen in which two or more drugs are alternated frequently during the treatment of a patient. The idea is that a rapidly changing environment impedes the adaptation of the bacteria to the antibiotics and consequently reduces the fraction of treatment failures due to resistance evolution [1].

The idea of alternating drugs was first studied as a strategy to prevent the spread of resistance across patients within a hospital (see Uecker, and Bonhoeffer [4] for a review of modelling studies). As such, it is implemented on a population level: the default drug used for empirical therapy – i.e., before identifying the specific disease-causing bacterium – in a hospital is cycled in time, but any single patient (ideally) receives just one drug throughout their treatment. While this hospital-wide cycling of drugs aims to reduce the spread of resistance between patients, sequential therapy during treatment of individual patients aims to minimise the probability of within-host resistance evolution. Usually, bacteria would be exposed to much more rapidly changing environments during sequential therapy than during hospital-wide drug rotations.

Laboratory experiments have demonstrated the superiority of fast sequential therapy over mono-therapy [5–8]. These studies showed that the treatment minimises the rate of adaptation and constrains the evolution of multi-drug resistance. It was further found that fast sequential therapy can lead to the clearance of bacterial populations under antibiotic concentrations for which the simultaneous administration of two drugs would fail [9], and that for certain drug pairs, a high switching rate correlates with a high extinction rate [8, 10].

Recent studies suggest that the efficiency of sequential therapy could potentially be further increased through the exploitation of evolutionary trade-offs such as collateral sensitivity [11, 12]. With collateral sensitivity, resistance to one drug increases the susceptibility to another one, which was first described by Szybalski, and Bryson [13]. This trade-off has been observed in different bacterial species, for instance in *E. coli* [11, 13–16], *P. aeruginosa* [17, 18], *S. faecalis* [12] and *S. aureus* [5, 19]. Identification of collateral sensitivity profiles for different drugs has indeed been successfully used to inform optimal treatment strategies, which outperform other sequential regimens and mono-therapy, in the lab [12].

In a laboratory setting, it is possible to switch abruptly from one drug to another. In a patient, in contrast, the antibiotic concentration decreases gradually due to pharmacokinetic processes such as metabolization or secretion of the drug. Doses of different drugs might then overlap such that drug-drug interactions can influence the dynamics. To be able to transfer results from the lab to the clinic, it is important to understand how pharmacokinetic processes influence the outcome of sequential therapy and the optimal treatment settings.

Mathematical models that take the pharmacokinetic and pharmacodynamic properties of drugs into account can simulate a patient-like environment and can help to close the gap between laboratory experiments and clinical applications. Such models have recently been applied to assess the benefits of collateral sensitivity for cycling strategies, showing that the benefits depend on the pharmacodynamic properties of the drugs and the drug dosing [20, 21]. However, these studies only compare a few cycling regimens and drug doses and, more importantly, do not include drug-drug interactions, which are one of the greatest differences between the lab and the patient. Moreover, they do not provide an explicit comparison to a ‘lab model’ with alternating constant drug concentrations.

In this article, we set up a lab model and a patient model. Our goal is two-fold – compare sequential therapy in the lab and in the patient and identify the optimal settings for patient treatment, depending on drug dosing, the pharmacodynamic drug characteristics, drug-drug interactions, and collateral sensitivity. This requires choosing a measure for the assessment of treatment strategies. We focus on dosing regimens in which the evolution of resistance slows down the decline of the bacterial population but ultimately does not prevent its extinction. We, therefore, choose the time until the sub-population sizes of all types have dropped below a given threshold as our primary measure. The analysis shows that the optimal frequency may indeed differ between sequential therapy in the lab and in the patient and that drug-drug interactions can have a strong effect on the treatment efficiency and should thus be taken into account when choosing drug pairs for therapy.

## 2 Methods

### 2.1 General Model

We consider a bacterial population of initial size *N*_0_ that undergoes antibiotic treatment following a certain regimen, which includes at most two drugs – drug A and drug B. The parameter *T* defines the time between two administrations. At each administration, either drug A or drug B is given at a dose *D_A_* or *D_B_*, and the drug sequence defines the treatment regimen (Figure 1A).

**Figure 1:**
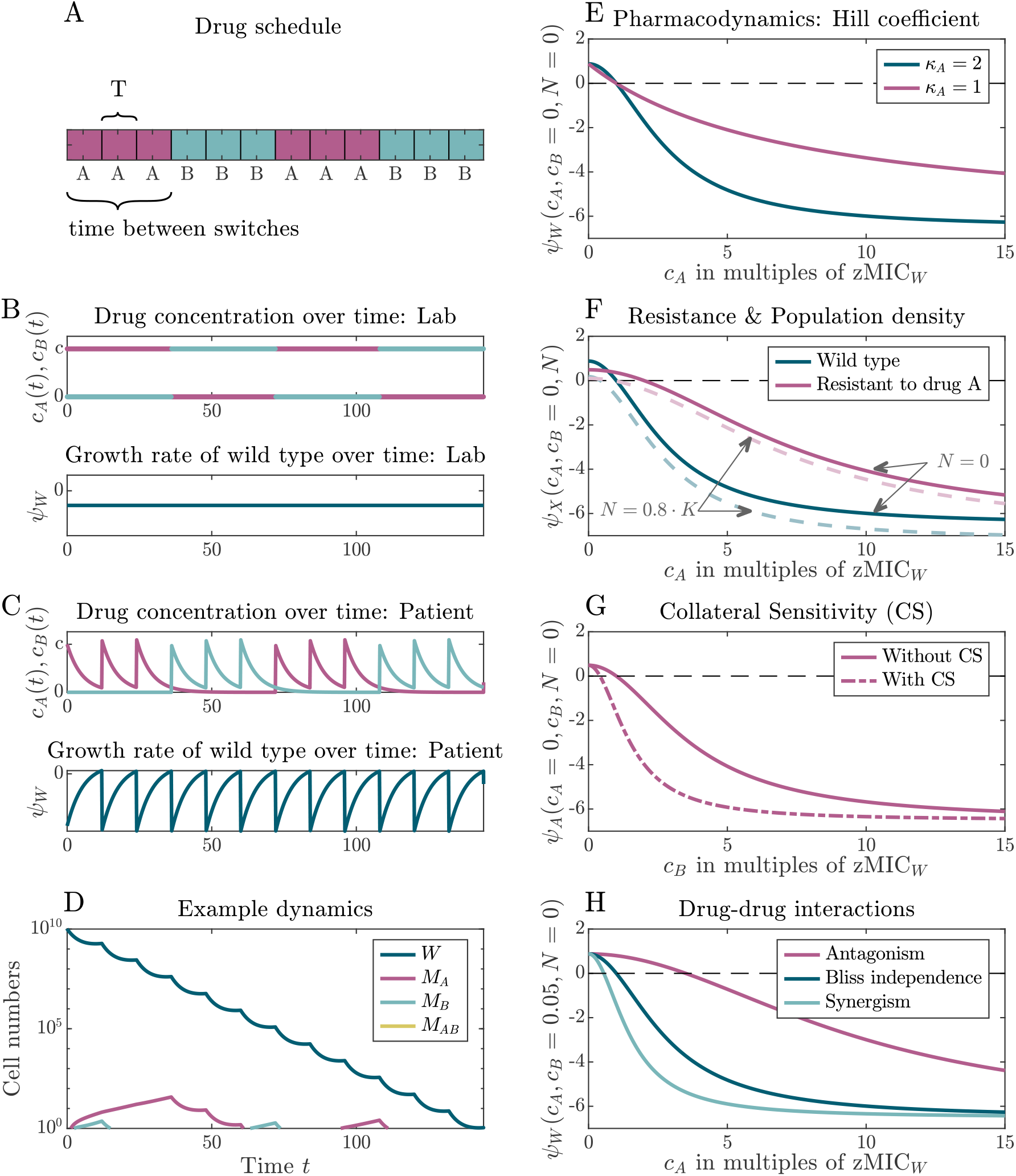
Illustration of the components of the PKPD model. **Panel A**: Example treatment schedule, in which the drug is switched after three administrations. **Panels B** and **C**: Drug concentration and growth rate of wild-type bacteria over time in the lab (Panel B) and in the patient (Panel C). **Panel D**: Dynamics of the bacterial strains for the treatment schedule displayed in Panel A. The y-axis starts at one cell. **Panels E-H**: Pharmacodynamic curves, illustrating several features. Panel E compares the curves for two different Hill coefficients *κ*; Panel F compares the curves for the wild-type and the single resistant strain at different cell densities; Panels G and H show the influence of collateral sensitivity and drug-drug interactions, respectively. (Parameter values differing from those in Table 1: 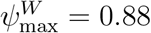, *β*_res_ = 2, *γ_A_* = *γ_B_* = *γ_AB_* = 0.4)

We consider four different types of cells: wild-type cells that are susceptible to both drugs, single mutants resistant to drug A or drug B, and double mutants that are resistant to both drugs. With probabilities *u_A_* or *u_B_*, a bacterium acquires a resistance mutation to the respective drug during replication.

**Table 1:**
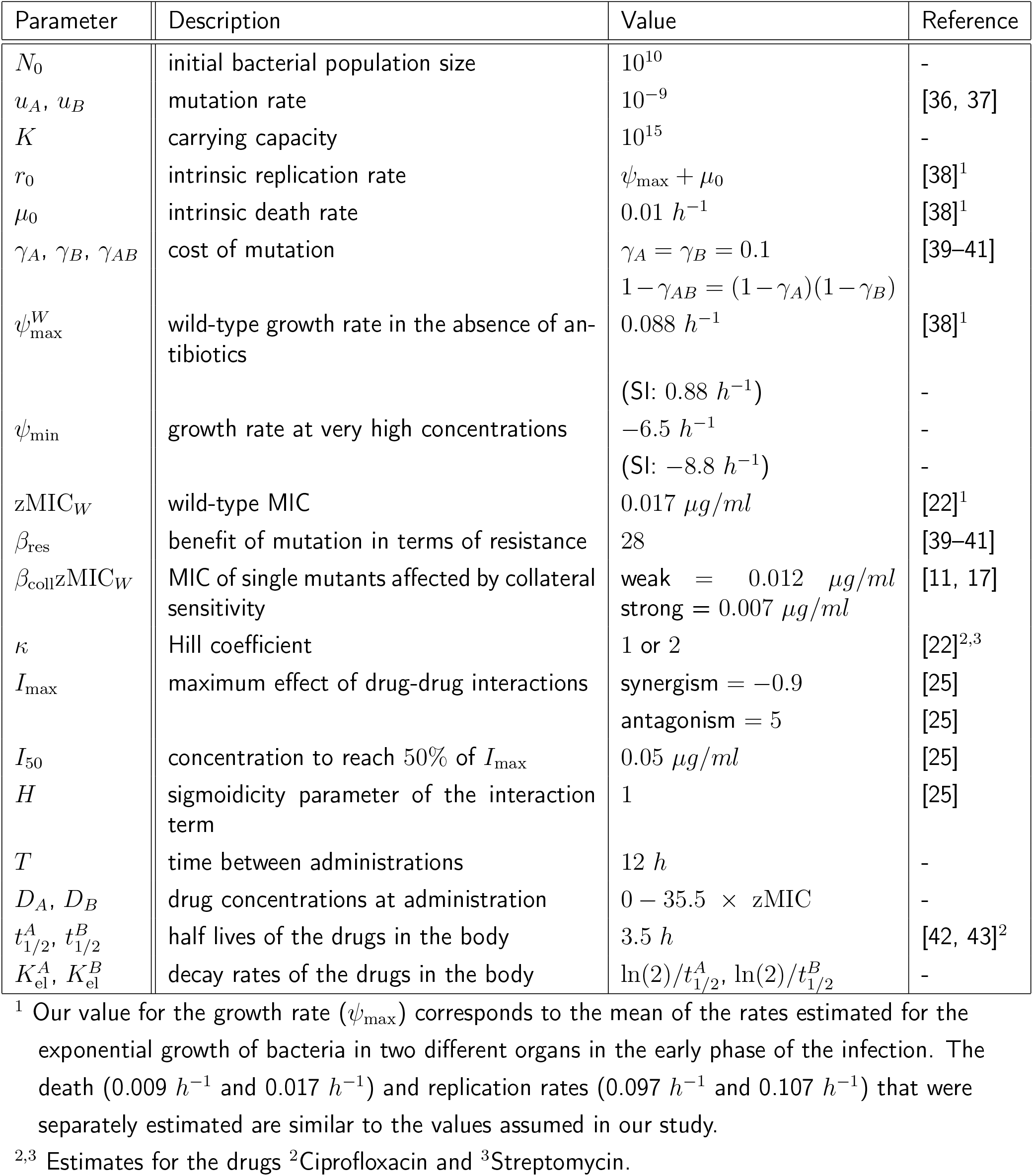
Table of all parameters and their values used to generate the results. When multiple values are given, these were varied during the analysis to observe the effect of a certain parameter on the treatment efficiency.

We assume that the drugs are strictly bactericidal, i.e. they increase the death rate of bacteria but do not affect replication. Replication of wild-type cells occurs with the density-dependent rate 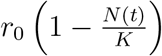, where *r*_0_ is the intrinsic replication rate, *N* (*t*) the total population size at time *t* and *K* the carrying capacity. Resistance entails a cost, reducing the maximum replication rates by factors (1 − *γ_A_*), (1 − *γ_B_*), and (1 − *γ_AB_*) for bacteria that are resistant to drug A, drug B, or to both drugs, respectively. The death rates are given by the sums of the intrinsic death rate *μ*_0_ and the antibiotic-induced kill rates *μ_X_* (*X* ∈ {*W, A, B, AB*}), which are type-specific and concentration-dependent and explained in detail in the next sections. The antibiotic concentrations of the two drugs, *c_A_*(*t*) and *c_B_*(*t*), change with time according to the treatment schedule and – for the patient – the pharmacokinetics of the drugs. We discuss the time course of *c_A_*(*t*) and *c_B_*(*t*) further below.

In serial transfer experiments, the bacterial population size undergoes a bottleneck before every drug administration. In the main text, we do not incorporate this dilution step into the lab model. The comparison between the patient and the lab thus reduces to a comparison between models with and without pharmacokinetics. Results for a more realistic lab model with bottlenecks are discussed in the supplement (SI section S3).

Putting everything together, the following ODE system describes the dynamics for each type, where *W* denotes the number of wild-type cells and *M_A_*, *M_B_*, and *M_AB_* the numbers of cells that are resistant to drug A, drug B, or to both drugs:

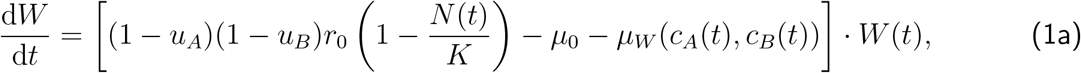

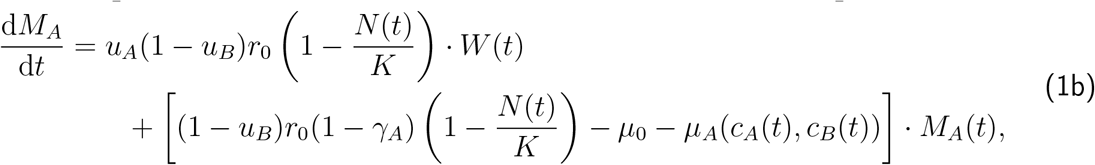

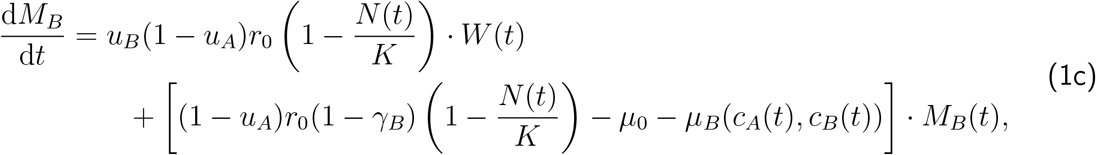

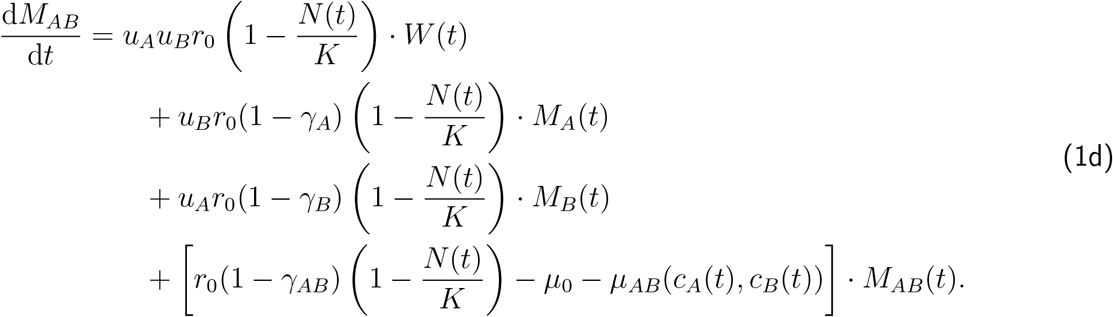

For most of our analysis, we assume that the population initially consists only of wild-type bacteria and consider the *de novo* evolution of resistance during treatment. We later test how the pre-existence of resistance affects our results.

### 2.2 The pharmacodynamics of the drugs

We next describe how we model antibiotic-induced killing. For notational simplicity, we drop the subscripts indicating the cell type and the drug for the moment. We start by describing the effect of a single drug that is present at concentration *c*. We denote by *Ψ*_max_ the growth rates of the respective type in the absence of antibiotics at low bacterial densities (e.g.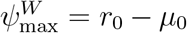 for the wild type and 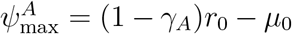 for the type resistant to drug A).

Following Regoes et al. [22], we describe the antibiotic-induced killing by the term

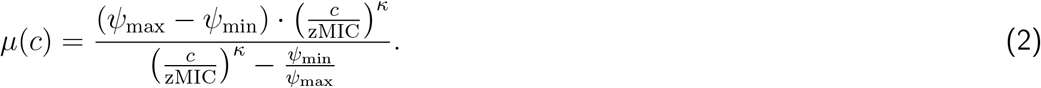

Antibiotic killing is thus independent of the bacterial density and the actual density-dependent replication rate of cells. The meaning of the parameters *Ψ*_min_, zMIC, and *κ* is best seen by considering the growth rate of bacteria in the absence of other cells

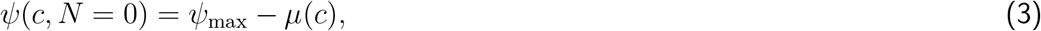

which is shown in Figure 1E: the parameter *Ψ*_min_ is the growth rate in the limit of very high antibiotic concentrations *c* (lim*_c→∞_Ψ*(*c*) = *Ψ*_min_). The parameter zMIC is the minimal inhibitory concentration (*Ψ*(zMIC) = 0). The so-called Hill coefficient *κ* regulates the steepness of the sigmoidal curve (compare the two curves in Figure 1E). Figure 1F shows the bacterial growth rate at non-zero population densities, where the parameters zMIC and *Ψ*_min_ loose this simple interpretation.

The pharmacodynamic parameters *Ψ*_min_, zMIC, and *κ* generally differ between the two drugs. Since we measure drug concentrations in multiples of the minimal inhibitory concentration of the wild type, the absolute minimal inhibitory concentrations with respect to the drugs do not matter, and we can choose the same for both without loss of generality; hence we set 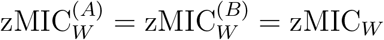. For simplicity, we also assume that *Ψ*_min_ is the same for both drugs.

We assume that – besides *Ψ*_max_ – only the minimal inhibitory concentration zMIC differs between the four cell types, while *Ψ*_min_ and *κ* are the same for all of them. This assumption conforms with experimental measurements of mutant dose-response curves [23, 24], which, however, does not mean that it is universally true. Bacteria that are resistant to a given drug can tolerate higher concentrations than the drug-sensitive wild type, i.e., their minimal inhibitory concentration with respect to that drug is higher. We model resistance by multiplying the wild-type parameter 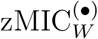 (where the • denotes the drug) by a factor *β*_res_ > 1 for the respective single mutant and the double mutant (Figure 1F). For example, the minimal inhibitory concentration of the mutant resistant to drug *A* with respect to drug *A* is 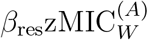. We also allow for collateral sensitivity. A single mutant displaying collateral sensitivity has a *lower* minimal inhibitory concentration than the wild type in the presence of the antibiotic to which it is susceptible. Similar as for resistance, we model this by a factor *β*_coll_ < 1 (Figure 1G). For example, the minimal inhibitory concentration of the mutant resistant to drug *A* with respect to drug *B* is 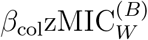. We assume that the double mutant is not affected by collateral sensitivity.

### 2.3 Combined effect of two drugs and drug-drug-interactions

If drugs are cycled rapidly, both drugs may be simultaneously present in the body. Interactions between the drugs can alter the individual effects of each drug. To account for this, we multiply the minimal inhibitory concentration of a drug with an interaction term *I* that depends on the concentration of the other drug [25]; see SI section S1.1. E.g., the wild-type MIC of drug *A* in the presence of drug *B* is 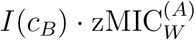. Following Wicha et al. [25], we model the interaction term by

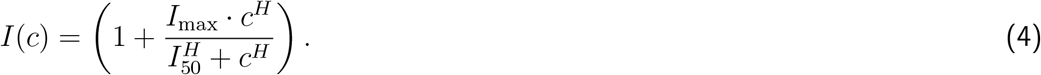

The variable *I*_max_ describes the maximal effect (lim*_c→∞_ I*(*c*) = 1 + *I*_max_). The sign of *I*_max_ determines whether the interaction is enhancing or inhibiting the effect of the focal drug. At concentration *c* = *I*_50_, we have *I* = 0.5*I*_max_. The steepness of the sigmoidal curve is given by the parameter *H*. For simplicity, we assume that effects are reciprocal (this holds for some interactions, but others are directional; for a review see [26]).

Finally, to obtain the combined effects of both drugs, we sum their individual kill rates. In the absence of drug-drug interactions, this corresponds to Bliss independence [27, 28]. For clarity, we explicitly give the full expression for the antibiotic-induced death rate of the wild type:

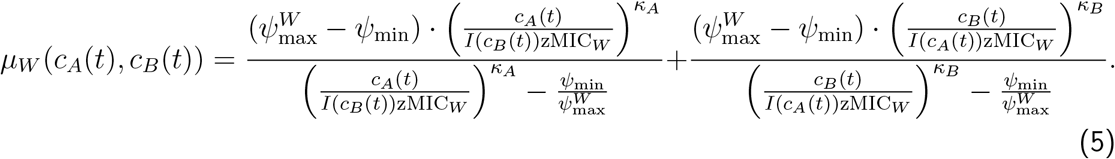

The growth rate is given by

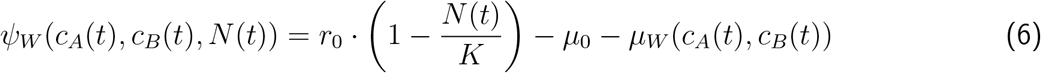

and illustrated in the presence and absence of drug-drug interactions in Figure 1H. The death and growth rates of the other types are obtained accordingly, using the respective maximal growth rates and minimal inhibitory concentrations.

With this choice of modelling drug-drug interactions, antagonistic interactions are hyper-antagonistic – also termed ‘suppressive’ – in some concentration range. I.e. wild-type bacteria grow *better* if the second drug is added than they would if the second drug were absent (*μ_W_* (*c_A_, c_B_*) < *μ_W_* (*c_A_*, 0) for certain concentrations *c_A_*, *c_B_*; see SI Figure S1A,E in SI section S1.2). However, they always grow worse than the mutant types (SI Figure S1B-D,G-H).

### 2.4 The pharmacokinetics of the drugs

We assume that in the lab, the antibiotic concentration stays constant at the applied dose *D_A_* or *D_B_* between administrations. This means that *c_A_*(*t*) is equal to *D_A_* whenever the schedule indicates a cycle of drug *A*, but is zero for the cycles of drug *B* (Figure 1B).

In the patient, in contrast, the drug concentration at the site of infection changes over time due to the pharmacokinetic processes of drug absorption, distribution, elimination, and metabolization [29]. To derive the functions *c_A_*(*t*) and *c_B_*(*t*), models in which the body is divided into different compartments can be used [30]. We here simply consider a one-compartment model. In the one-compartment model, the entire dose *D_A_* (or *D_B_*) is fully available after administration and eliminated from the body at rates 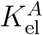 (or 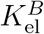). The rates 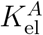 and 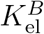 can be estimated from the drug’s half lives 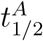 and 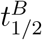. Let us denote by 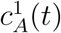 the exponential decline in the concentration of drug *A* after one administration at time *t* = 0 (equivalent for drug *B*):

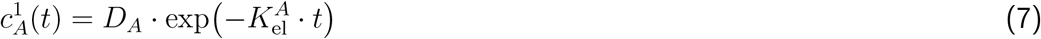

During the time course of the treatment, both drugs are administered multiple times. *c_A_*(*t*) and *c_B_*(*t*) are then given by the sum of time-shifted functions 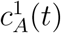 and 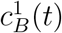 based on the defined treatment regimen and the pharmacokinetic parameters for each drug (Figure 1C, upper graph). Together with the pharmacodynamics described in the previous section, this leads to time-dependent growth rates (Figure 1C, lower graph). For all numerical results, we assume that the administered doses and the decay rates of the drugs are the same for both drugs (*D_A_* = *D_B_* = *D*, 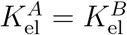).

### 2.5 Implementation

We solved the deterministic ODE system Eq. (1) numerically in MATLAB (version 2020a/2020b) using *ode45* [31]. Table 1 provides a list of all parameter values. Example dynamics are shown in Figure 1D.

Stochastic effects can become important when the population size gets small. Therefore, we additionally set up a hybrid model that accounts for stochasticity when the sub-population size for a type drops below a certain threshold (see SI section S1.3). Below this threshold (here set to 1000), we simulate the cell division and death by the respective rates, applying the thinning algorithm introduced by Lewis, and Shedler [32]. This algorithm is based on the Gillespie algorithm [33, 34] but accounts for time-dependent rates. When a type exceeds the threshold, the cell numbers are calculated with the deterministic model described above; the only exception are mutations, which are always simulated as stochastic events. Simulations with this hybrid approach (described in Kiehl et al. [35]) are faster than fully stochastic simulations. However, they are not fast enough to conduct the same analysis we are able to do with the deterministic model. In SI section S1.4, we compare results of the two model implementations for a small set of treatments. Overall, the results align well. However, with increasing concentrations, the agreement gets less good. We therefore support all results from the deterministic model with semi-stochastic simulations throughout the main text. The hybrid model moreover allows assessing how robust the performance of a treatment regimen is with respect to stochastic effects, which is discussed in the supplementary information section S1.5.

### 2.6 Assessing the performance of a treatment regimen

To compare different treatment regimens with each other, we need to define a measure for the performance of a treatment. We focus on concentrations at which, for a given treatment, the appearance of resistant strains does not completely prevent suppression of the bacterial population but may slow down its decline. Importantly, as we will see, this does not imply that we apply concentrations above the MIC of the single resistant strains. As the main model implementation is deterministic, eradication of the bacteria is not possible. We therefore use the time until each sub-population size has been reduced to a threshold *N_c_*, which we set to one, as a measure of treatment efficiency. As a second proxy for treatment efficiency we consider the minimal concentration for which each sub-population drops below *N_c_* within the considered time frame of 80 drug administrations (40 days). We call this concentration the first effective concentration. For the semi-stochastic simulations, we only show results in a range of doses in which treatment is successful in all replicate runs; the lowest concentration, for which results are shown, is further limited by the run time of the semi-stochastic simulations and can thus not be interpreted as the first effective concentration.

## 3 Results

To identify potential differences between the lab and the patient model and to determine the optimal settings for sequential therapy, we will compare in this section the efficiency of different treatments, varying in cycling frequency and drug characteristics. We consider a broad range of drug concentrations so that we are able to determine the effect of the treatment settings for low and high drug concentrations.

In laboratory experiments, bacteria are usually provided with optimal growth conditions and replicate quickly. *In vivo*, in some infections, replication is fast as well, while in others bacteria replicate very slowly (see [38] or [44] for studies in mice). We therefore study treatment of both slowly and rapidly replicating bacteria. It turns out that some results are sensitive to the rate of cell division and that pharmacokinetic processes alter conclusions for slowly replicating but not for rapidly replicating bacteria. In the main text, we focus on results for slowly replicating bacteria (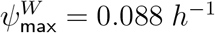) and point out differences to rapidly replicating bacteria 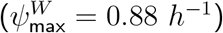 throughout. We additionally summarize all results in Table 2 at the end of the results section. The full analysis for rapidly dividing cells, including a detailed comparison of the dynamics, can be found in SI section S3.

**Table 2:** Summary of main results for the treatment of slowly replicating and rapidly replicating bacteria in the lab and in the patient.

|  | Replication | Results | Figure |
| --- | --- | --- | --- |
| Optimal cycling frequency | Slow | The most rapid cycling is always the best strategy or at least as good as slower cycling regimens. | 3 |
|  | Rapid | The most rapid cycling is always the best strategy or at least as good as slower cycling regimens | S17 |
| Collateral sensitivity (CS) | Slow | CS has mainly an effect on the treatment efficiency of slow cycling regimens, the effect decreases with increasing concentrations (no effect at very high concentrations). | S10 |
|  | Rapid | In the absence of bottlenecks, CS has mainly an effect on the efficiency of fast cycling regimens at low concentrations but the effect is weak. In the presence of bottlenecks, CS might also slightly increase the treatment efficiency of slow cycling regimens at low concentrations. | S23 |

|  | Replication | Results | Figure |
| --- | --- | --- | --- |
| Optimal cycling frequency | Slow | The most rapid cycling is not always optimal, slower cycling can be optimal if $\kappa_A = \kappa_B = 2$ or under antagonistic drug-drug-interactions. | 3, 5 |
|  | Rapid | The most rapid cycling is always the best strategy or at least as good as slower cycling regimens. | S17 |
| Drug-drug interactions | Slow | Drug-drug-interactions have mainly an effect on the treatment efficiency of fast cycling regimens, the effect slightly decreases with increasing concentration. | 4 |
|  | Rapid | Drug-drug-interactions have mainly an effect on the treatment efficiency of fast cycling regimens, but the effect is less strong. | S28 |
| Collateral sensitivity (CS) | Slow | CS has mainly an effect on the treatment efficiency of slow cycling regimens, the effect vanishes at very high concentrations. CS has an effect on fast cycling regimens when resistance is pre-exists. | 4, 6 |
|  | Rapid | CS has mainly an effect on the treatment efficiency of fast cycling regimens at low concentrations, but the effect is weak. (Pre-existence of resistance was not considered.) | S23 |

In the first part of the analysis, we evaluate the efficiency of the sequential regimens in comparison to mono-therapy.

We then explore in the second part the influence of the pharmacodynamics of the drugs. Our focus lies on the Hill coefficient *κ*. We briefly discuss the effect of the pharmacodynamic parameter *Ψ*_min_ in SI section S2.1.

In the third part, we analyse the effect of collateral sensitivity and drug-drug interactions on the treatment efficiency. For the latter, we assume either synergism, in which both drugs potentiate each other’s effect, or antagonism, in which both drugs inhibit each other’s effect.

In the last part, we briefly discuss results for a heterogeneous initial population, where resistance pre-exists prior to treatment.

In all figures, discrete data points are connected by lines for easier readability.

### 3.1 Sequential therapy performs better than mono-therapy

Figure 2 shows the first effective concentration for a large range of treatments, highlighting the need for a higher concentration to achieve a treatment success under mono-therapy than under sequential therapy (blue lines). Importantly, sequential therapy is effective well below the MIC of the single mutants (dashed grey line). The reason for the reduced first effective concentration under sequential therapy could, in principle, either be a more efficient clearance of the wild type or a better suppression of resistant strains under drug cycling. To disentangle the two, we also considered the first effective concentration if resistance evolution were impossible, i.e. *u_A_* = *u_B_* = 0 (yellow lines). In that case, the first effective concentration is virtually the same for mono-therapy and for all sequential regimens, showing that sequential therapy is more efficient than mono-therapy through more effectively controlling the growth of the resistant types (but we will see further below that the cycling frequency can also have an effect on the decline of the wild-type population). To give an upper bound for the concentration needed for successful treatment, we furthermore included the results for a population that is entirely composed of double mutants (lilac lines). A comparison of Panels A and B shows that successful treatment requires lower concentrations in the lab than in the patient, which is expected given that the concentration decreases over time in the patient but not in the lab.

**Figure 2:**
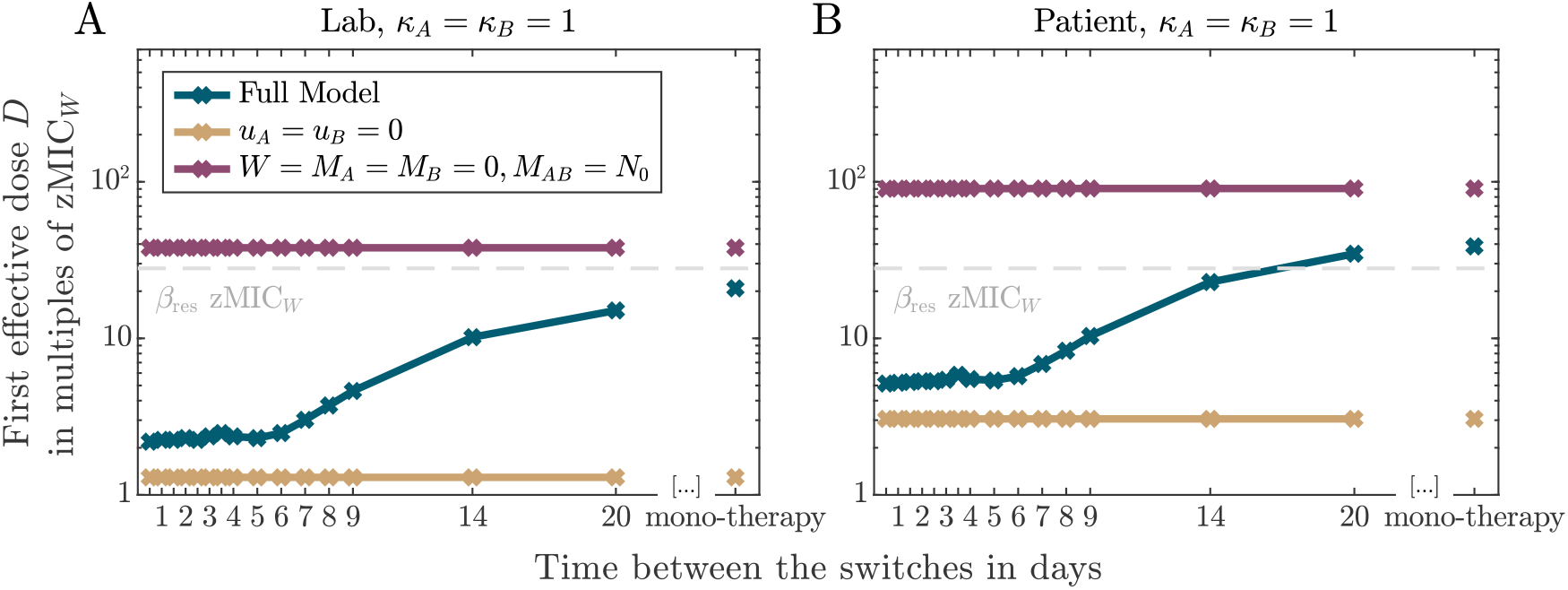
First effective concentration for a large range of sequential regimens in comparison to mono-therapy for the lab and patient environment. The blue curves shows the results for the model as described in the methods section; the yellow curve shows results for a fully susceptible cell population in which resistance evolution is impossible (*u_A_* = *u_B_* = 0); the lilac curve shows results for treatment of a population of double resistant cells. The grey dashed line marks the MIC of the single mutants.

### 3.2 The optimal cycling frequency for patient treatment depends on the Hill coefficients of the drugs and the antibiotic dose

In this section, we focus on the effect of the drugs’ Hill coefficients *κ_A_* and *κ_B_*, which reflect the steepness of the dose-response curves on the treatment. For simplicity, we consider treatments with two drugs that have the same Hill coefficient (either *κ_A_* = *κ_B_* = 1 or *κ_A_* = *κ_B_* = 2). Example simulations for drug pairs with different Hill coefficients (*κ_A_* = 1 and *κ_B_* = 2 or the other way round) can be found in SI section S2.2.

Figure 3 compares the different treatment regimens for the two different drug pairs (*κ_A_* = *κ_B_* = 1 and *κ_A_* = *κ_B_* = 2) in the lab and in the patient. Considering the semi-stochastic simulations, we see that differences between the strategies are more pronounced in the lab than in the patient over most of the parameter range. In most cases, the treatment efficiency decreases with the time between drug switches, with several rapid regimens performing equally good. In particular, in the lab model, rapid cycling minimizes the time to extinction across the entire concentration gradient for both drug pairs. In the patient, the picture is more nuanced. For the drug pair with Hill coefficients *κ_A_* = *κ_B_* = 2, the optimal cycling frequency depends on the drug concentration. At low drug concentrations, a little bit slower cycling frequencies are slightly better than very high cycling frequencies (see Figure 3G and H). Why could this be the case? When a new dose is administered, there is still some residual antibiotic present in the body from the previous administration. One of the differences between the various cycling regimens lies in the number of overlaps between doses of different drugs and doses of the same drug. For the fastest cycling regimen, overlaps occur only between doses from different drugs. In SI section S2.3, we argue that for drug pairs with a ‘large’ Hill coefficient and low antibiotic concentrations, dose overlaps from the same drug lead to higher killing than dose overlaps from different drugs for all cell types except for the one that is resistant to this drug. This type is then managed by administrations of the other drug, making intermediate cycling frequencies overall optimal. For rapidly replicating cells, the concentration for which same-drug overlaps provide an advantage are lower than the first effective concentration. In this case, the most rapid cycling is always optimal, even for drugs with ‘large’ Hill coefficients (see Figure S17).

**Figure 3:**
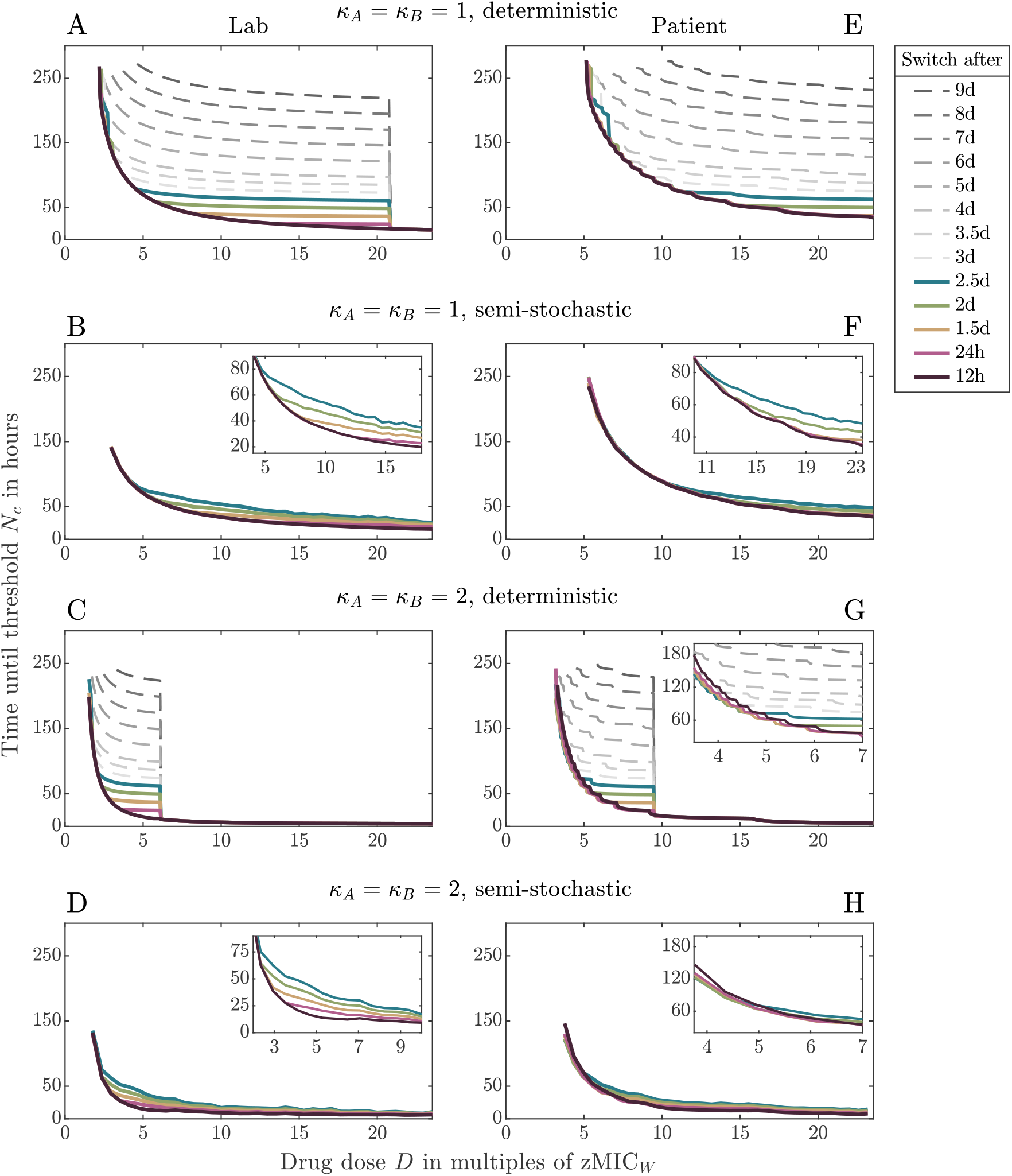
Treatment efficiency of di fferent reg imens, cyc ling dru g pai rs wit h eit her *κ_A_* = *κ_B_* = 1 (upper four panels) or *κ_A_* = *κ_B_* = 2 (lower four panels) in the lab (left column) or in the patient (right column). Each plot compares the time by which all sub-population sizes reach the threshold *N_c_* for different t reatments a cross a wide range of concentrations. Panels A,C,E,G show results obtained from the deterministic ODE model Eq. (1); Panels B,D,F,H show results from simulations based on the semi-stochastic hybrid model. The most rapid cycling regimen is always optimal in the lab. In the patient, slightly slower cycling performs better at low drug concentrations for the drug pair with *κ_A_* = *κ_B_* = 2 (see Panels G and H).

The deterministic ODE model correctly captures the behaviour in all scenarios but overestimates differences between strategies for low/intermediate concentrations and underestimates them for high concentrations. In particular, in the deterministic model, all strategies (except for the 12*h* treatment) are equally good from a certain concentration on (see SI section S1.4 for details).

### 3.3 Collateral sensitivity influences the efficiency of slow cycling regimens and drug-drug interactions influence the efficiency of fast cycling regimens

In this third part, we describe how collateral sensitivity and drug-drug interactions influence the efficiency of a given cycling regimen and the optimal treatment choice. In all cases, we consider these characteristics to be reciprocal, e.g. collateral sensitivity in both single mutants and reciprocal drug-drug interactions. Results for collateral resistance (i.e. *β*_col_ > 1) are discussed in SI section S2.4.

As shown in Figure 4A and D, for slowly replicating bacteria, collateral sensitivity does not increase the treatment efficiency when the drugs are switched rapidly. As the cycling frequency decreases (Panels B, C and E), however, collateral sensitivity becomes important (see SI section S2.5 for results for the lab model). For drug-drug interactions, we observe the opposite – the efficiency of fast cycling is strongly influenced by the interactions, whereas slow cycling is not (Figure 4F-J). Antagonism leads thereby to a decrease and synergism to an increase in the treatment efficiency. The strong effect of drug-drug interactions on the performance of rapid but not slow cycling regimens is intuitive since rapid cycling leads to more periods of drug overlap during which the interaction becomes relevant. The results for the effects of collateral sensitivity are maybe less intuitive. One may think that collateral sensitivity should be more influential under fast than under slow cycling as well, as the trade-off can be exploited on more time points. The reason that this is not observed lies in the sizes of the resistant sub-populations during the treatments. The effect of collateral sensitivity becomes visible only when the affected sub-populations (i.e. the single resistant types) are sufficiently large. Rapid cycling is able to better suppress the resistant sub-populations than slow cycling, even in the absence of collateral sensitivity. Similarly, since the resistant sub-populations grow more slowly under high drug concentrations, the effect of collateral sensitivity becomes less visible with increasing concentration, as also observed by Aulin et al. [21].

**Figure 4:**
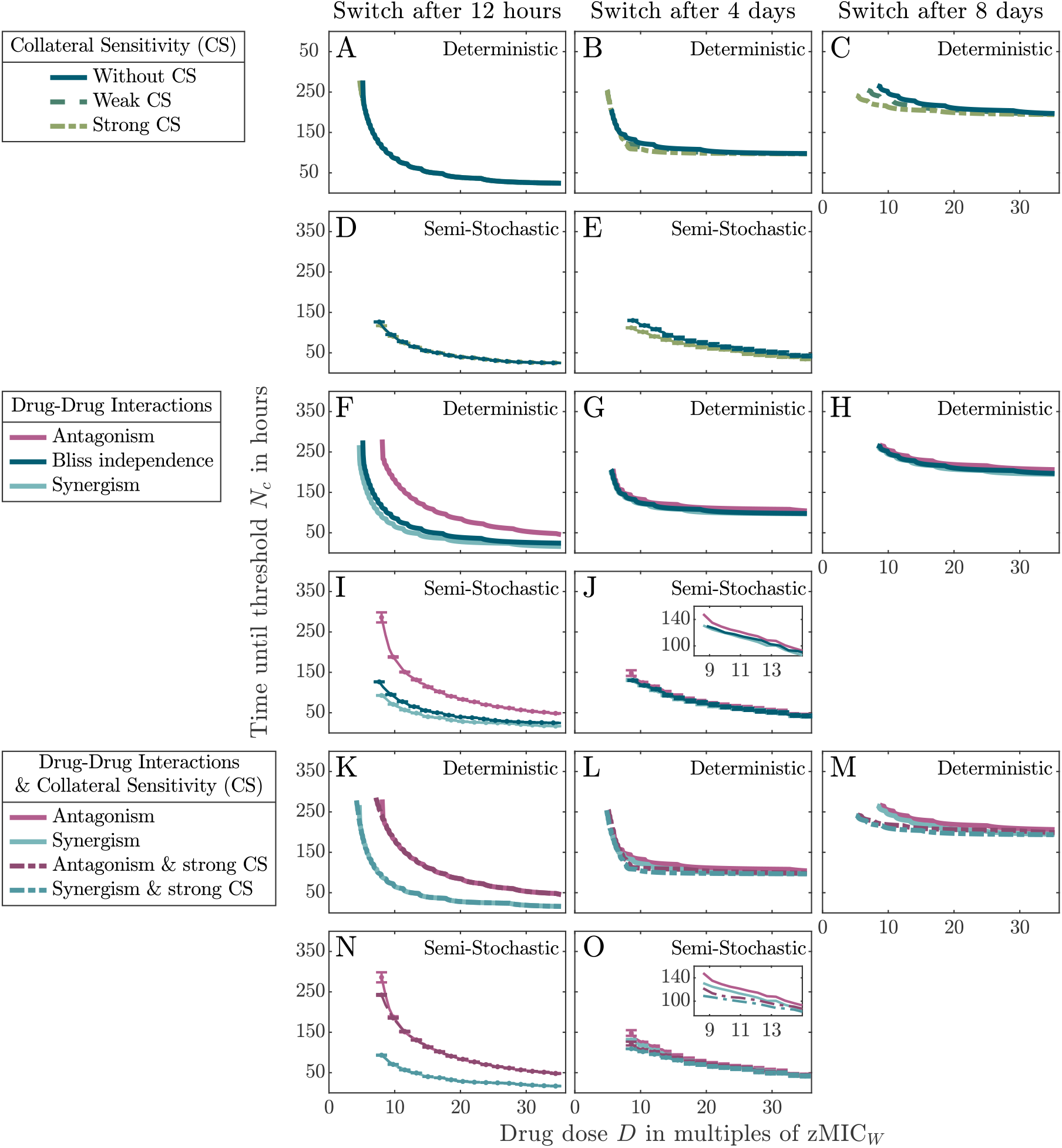
Treatment efficiencies under different drug characteristics for three different cycling regimens with *κ_A_* = *κ_B_* = 1. **Panel A-E**: Effect of collateral sensitivity, comparing efficiencies of drug pairs with weak and with strong collateral sensitivity of resistant mutants to the efficiency of a drug pair without collateral sensitivity. The influence of collateral sensitivity on the treatment efficiency increases as the cycling frequency decreases. **Panels F-J**: Effect of synergistic and antagonistic drug-drug interactions. Drug-drug interactions are most relevant in fast cycling regimens. **Panels K-O**: Combined influence of collateral sensitivity and drug-drug interactions. For slow cycling, a drug pair with antagonistic drug-drug interactions and collateral sensitivity is a better choice than a drug pair with synergistic drug-drug interactions but without collateral sensitivity. Panels A-C, F-H, K-M show results from the deterministic ODE21model; Panels D, E, I, J, N, O show results from the semi-stochastic hybrid model. Semi-stochastic simulations were too slow to simulate the regimen in which drugs are switched every 8 days.

If cells divide rapidly, resistant sub-populations can become large even if drugs are cycled fast. Collateral sensitivity therefore has an effect – albeit a very weak one – for rapid regimens at low drug concentrations both in the lab and the patient (Figures S23, S24A, and S25A). In contrast, collateral sensitivity effects are negligible for slow regimens (unless the population is subject to repeated bottlenecks, where at least the deterministic ODE model predicts a difference) because at the high concentrations required for treatment success, growth rates with and without collateral sensitivity are similar.

Until now, we have studied treatment efficiencies for drug pairs that either display drug-drug interactions or collateral sensitivity of resistant types but not both. Collateral sensitivity has been found for drug pairs with synergistic and for drug pairs with antagonistic interactions [45], and the presence of both, drug-drug-interaction and collateral sensitivity, can strongly alter the speed of adaptation compared to cases in which only one characteristic is present, as was observed in models for combination treatment (see Figure 4 in [46]). We show treatment efficiencies for such drug pairs in Figure 4K-O. Our analysis shows that if drugs are cycled slowly, an antagonistic drug pair with collateral sensitivity is a better option than a synergistic drug pair without collateral sensitivity (best visible in Panel M and in the inset of Panel O). The results in Figure 4 show that collateral sensitivity and drug-drug interactions influence the performance of some regimens more than that of others. Figure 5 compares the treatment efficiencies across cycling regimens for two selected scenarios in more detail (see SI section S2.6 for more scenarios). In line with the previous observations, we see that the characteristics of the drug pair can influence the relative performances of the cycling strategies. This is most prominently seen for the antagonistic drug pair, where very rapid cycling becomes a poor choice when treating slowly replicating bacteria. For rapidly replicating bacteria, in contrast, rapid cycling remains the optimal choice since in that case, the effect of drug-drug interactions is weak compared to the effect of the cycling frequency itself (Figure S30).

**Figure 5:**
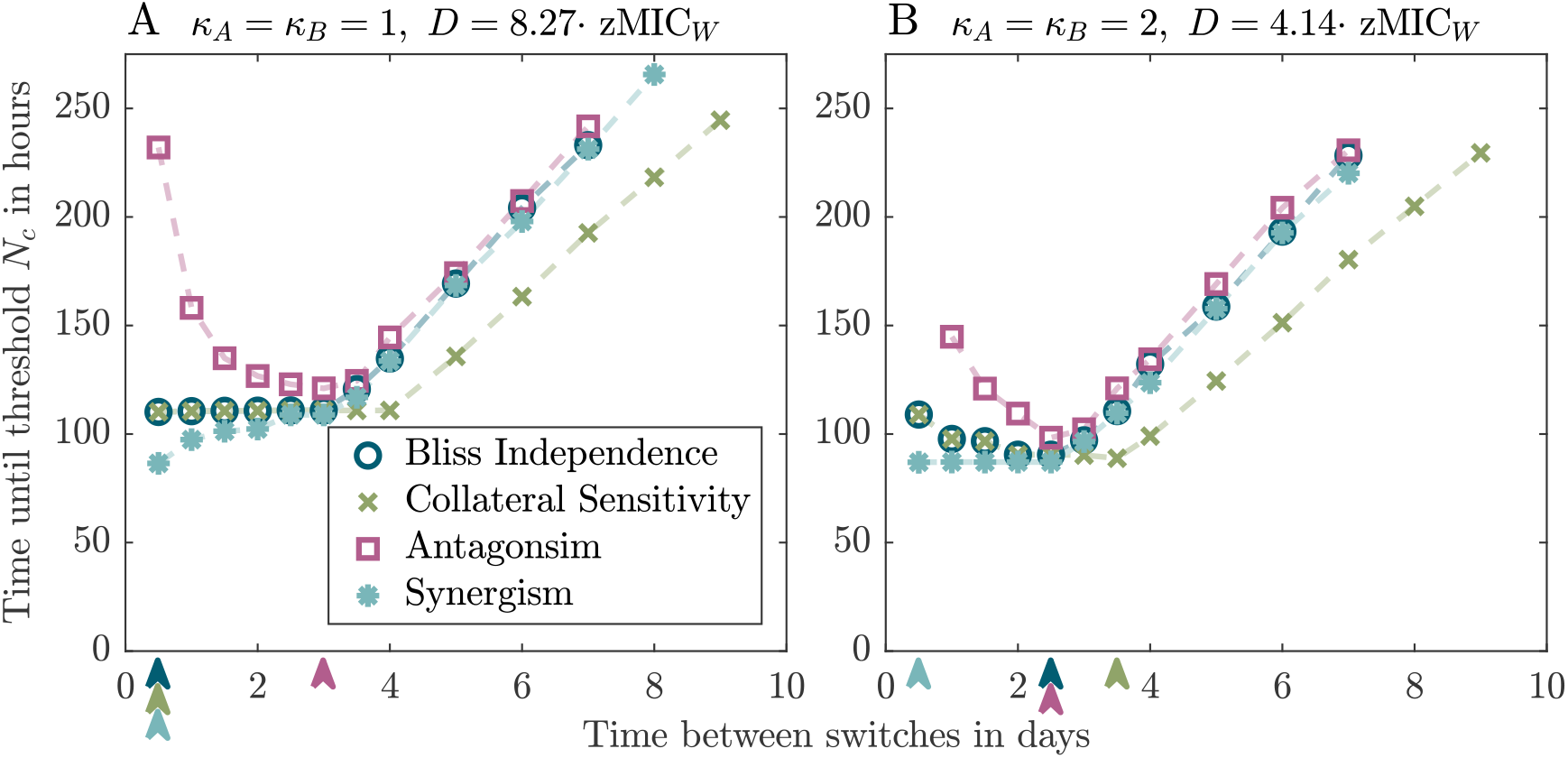
Comparison of treatment efficiencies for drug pairs with different drug characteristics at selected concentrations for *κ_A_* = *κ_B_* = 1 in Panel A and *κ_A_* = *κ_B_* = 2 in Panel B. The arrows indicate the cycling frequency that is optimal or at least as good as slower regimens for the respective characteristic. The figure shows results from the deterministic ODE model.

### 3.4 The pre-existence of single mutants decreases the treatment efficiency and makes collateral sensitivity beneficial even for rapid cycling

Up to now, we assumed that the initial population consists entirely of susceptible wild-type bacteria. However, resistant types may already be present prior to treatment. Here, we finally investigate the treatment efficiency for a heterogeneous population. To derive the number of pre-existing resistant cells, we simulated growth of a bacterial population from a single wild-type cell up to 10^10^ cells in the absence of antibiotics (i.e. we numerically solved the ODE system Eq. (1), setting antibiotic-induced killing to zero). By the time the population had reached 10^10^ cells, approximately 92 cells were resistant to drug *A* and drug *B* respectively; the double resistant type was absent (< 1 cell). To also investigate the effect of higher initial levels of resistance, we additionally consider a scenario with *M_A_*(0) = *M_B_*(0) = 1000 and *M_AB_* = 0.

As expected, the treatment efficiency is lower if resistance pre-exists, which especially affects slow regimens and for antagonism also very rapid regimens (SI Figure S13). Importantly, with increasing numbers of pre-existing mutants, collateral sensitivity becomes effective at low (but not at high) concentrations, even for rapid cycling (Figure 6). This effect is, however, much more pronounced for a drug pair with *κ_A_* = *κ_B_* = 1 than for a drug pair with *κ_A_* = *κ_B_* = 2, where collateral sensitivity becomes effective only over an extremely narrow range of the concentrations that we consider.

**Figure 6:**
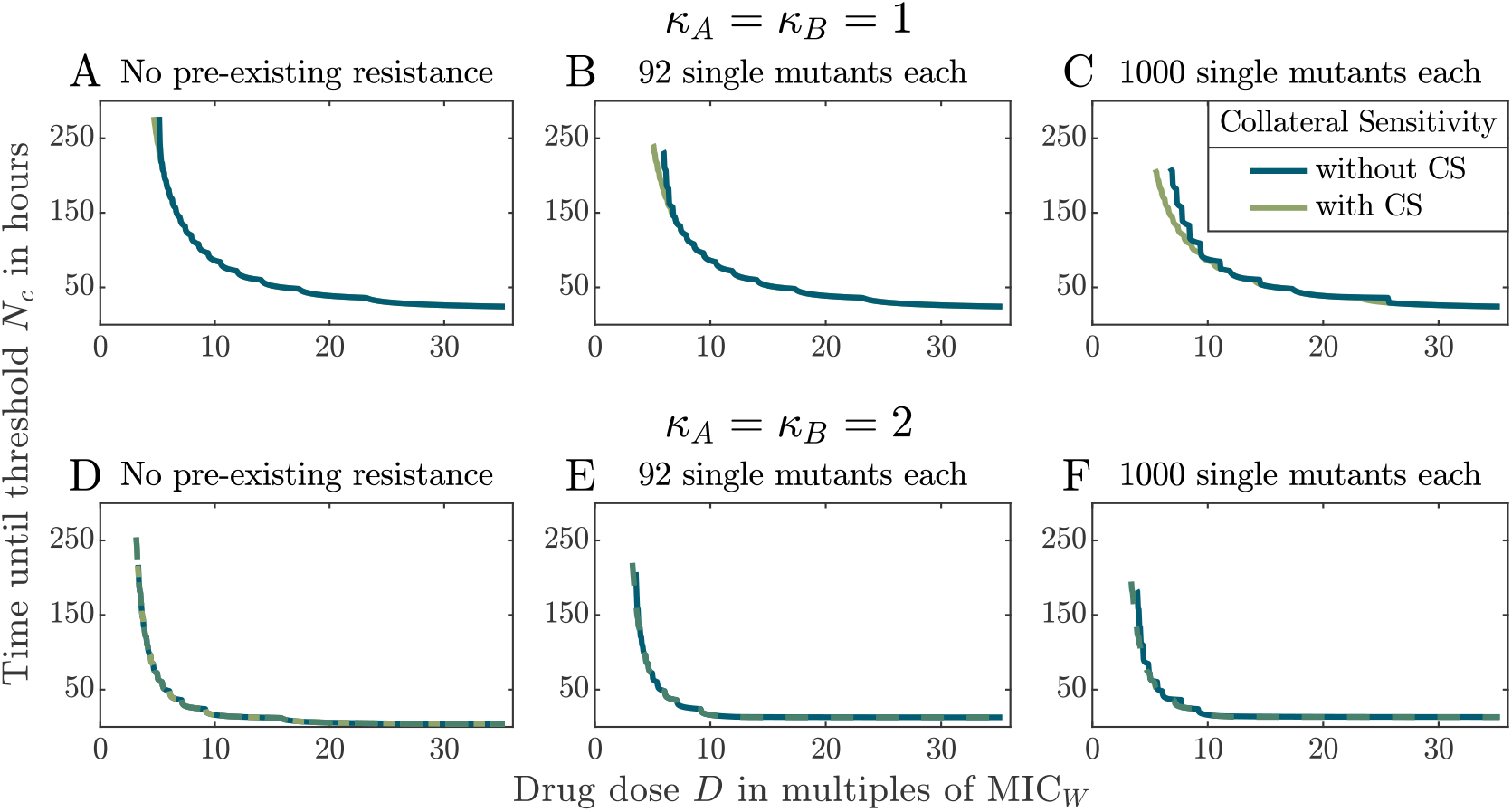
Comparison of treatment efficiencies for drugs pairs with and without collateral sensitivity for different starting populations in the columns, varying in the pre-existence of single resistant types. The graphs show the time until all sub-population sizes have been suppressed below a threshold size *N_c_*. Panel A-C show the treatment for drugs with *κ_A_* = *κ_B_* = 1 and Panel D-F for drugs with *κ_A_* = *κ_B_* = 2. Panel A and D displays the comparison for a starting population without the pre-existence of resistance. In Panel B and E or C and F the starting population includes 92 or 1000 single resistant bacteria of each type respectively.

## 4 Discussion

Which factors affect the efficiency and the optimal cycling frequency of sequential therapy? And is the optimal cycling frequency the same in the lab and in the patient? To answer these questions, we set up a pharmacokinetic-pharmacodynamic patient model and a lab model for comparison and simulated treatments for a set of example drug pairs. To make sure that our main conclusions are robust to stochasticity, we performed semi-stochastic simulations in addition to solving a deterministic ODE system.

### Comparison between the patient and the lab

One key difference between the lab and the patient are the pharmacokinetic processes that occur in the patient, which is the focus of the present study. Especially, unlike in the test tube, antibiotics from the previous administration may still persist in the body at the time of the next administration. For slow cycling, mostly doses of the same antibiotic overlap; for rapid cycling, most overlaps involve doses from different drugs. We find that conclusions from the lab model (with or without bottlenecks) mostly carry over to the treatment of rapidly dividing bacteria but not necessarily to the treatment of slowly dividing bacteria in the patient.

We find that in the lab model, the most rapid cycling performs either best or equally good as slower regimens, independent of the drug concentration, the replication rate of bacteria, and the presence or absence of bottlenecks. This finding is in line with the correlation between the cycling frequency and the extinction rate of bacterial populations observed in evolution experiments by Batra et al. [8] and Roemhild et al. [10]. For the patient, it holds true as well if the infecting bacteria have a high cell division rate. For treatment of infections with slowly dividing bacteria, by contrast, slightly slower cycling can sometimes lead to a higher treatment efficiency. As we will discuss in more detail below, very rapid cycling can in that case be an inferior strategy when (1) the pharmacodynamics curve is steep and drug concentrations low or if (2) the drugs interact antagonistically. We did not observe that collateral sensitivity ever made rapid regimens perform worse than slower regimes, but this could, in principle, happen in other parts of the parameter space.

### The influence of the Hill coefficients of the drugs on treatment efficiency and the optimal cycling frequency

How the steepness of the pharmacodynamic function influences treatment success and antibiotic resistance evolution has recently started to attract attention in modelling studies [47, 48]. A steeper pharmacodynamic function increases the mutant selection concentration, i.e. the concentration above which a resistant mutant has a selective advantage over the susceptible wild type [47, 48]. It entails lower selection for resistance at low concentrations and a larger kill rate at high concentrations [48]. In a PKPD model for mono-therapy, considering treatment of rapidly replicating bacteria, Yu et al. [48] found that treatment with a drug having a large Hill coefficient (in their case 5) leads to a lower probability of resistance and less treatment failure than treatment with a drug having a low Hill coefficient (in their case 1.5). Similarly, Aulin et al. [21] find that with slow drug cycling, drug pairs with large Hill coefficients (in their case 3) perform better than drug pairs with low Hill coefficients (in their case 0.5). For rapid cycling, in contrast, Udekwu, and Weiss [20] and Aulin et al. [21] find that drug pairs with lower Hill coefficients (in their cases 1 and 0.5 respectively) can be superior to drug pairs with larger Hill coefficients (in their case 3) in terms of delaying the rise of double resistance to large numbers [20] or in reducing the probability of resistance for rapid cycling [21].

For treatment of slowly replicating bacteria, we find that, irrespective of the cycling frequency, lower drug doses are sufficient for successful treatment and suppression of the bacterial population is faster for *κ_A_* = *κ_B_* = 2 than for *κ_A_* = *κ_B_* = 1, both in the lab and in the patient. For rapidly replicating bacteria, our results are more complex, and the first effective concentration is in many cases lower for the drug pair with *κ_A_* = *κ_B_* = 1. Essentially, the outcome is determined by the two opposite effects of the Hill coefficient below and above the MIC of bacteria: above their MIC, bacteria are killed more efficiently if the Hill coefficient is high, but below their MIC, they grow less if the Hill coefficient is low. Besides differences in parameters such as the population-wide mutation rate, one other reason why our results for treatment of rapidly dividing bacteria are not straightforward to compare to other studies is that we focus on the time to reduce all sub-population sizes to a given threshold as a measure of treatment quality.

Overall, the results across studies confirm that the Hill coefficient is an important parameter, the consequences of which are not always trivial. To complement the picture provided by our study, we performed a supplementary analysis in which we considered the number of double mutants at the end of the treatment for low drug doses and rapid drug cycling for slowly replicating bacteria (SI section S2.7). We find that resistance evolves over a smaller range of doses for *κ_A_* = *κ_B_* = 2 than for *κ_A_* = *κ_B_* = 1; especially, in line with the above observations for a single drug, the minimum dose above which resistance evolution occurs is higher. For a small range of doses, however, the final size of the double resistant sub-population is higher for the drug pair with the larger Hill coefficient.

In terms of the optimal cycling frequency, we find that the most rapid cycling regimen is always as least as good as slower regimens for *κ_A_* = *κ_B_* = 1 (unless drugs interact antagonistically), while slightly slower regimens can sometimes be preferable if the Hill coefficient is large (*κ_A_* = *κ_B_* = 2) and the bacteria are replicating slowly. Unlike us, Aulin et al. [21] find even for rapidly replicating bacteria that the optimal cycling frequency depends on the drugs’ Hill coefficient: In the absence of collateral sensitivity, the most rapid cycling regimen (1-day cycling) minimises resistance evolution for a low Hill coefficient, while for a large Hill coefficient, a single drug switch proves to be optimal (3-day cycling, which is intermediate between the two, performs worst).

### The influence of collateral sensitivity

It has been speculated that rapid cycling regimens could profit from exploiting collateral sensitivity effects [8]. In contrast with this idea, we found that if bacteria divide slowly and resistance evolves *de novo*, collateral sensitivity increases the treatment efficiency only for regimens with slow switching rates. The reason is that under rapid cycling, resistant sub-populations never become large enough for collateral sensitivity to have a visible effect. If single mutants pre-exist at sufficiently high levels at the start of treatment, however, collateral sensitivity can show an effect even for rapid cycling. If the cell division rate is high, the efficiency of rapid cycling regimens (but not of slow ones) is indeed improved by collateral sensitivity at low concentrations. However, the effect is very small. Throughout our study, we focus on drug concentrations that are high enough to ultimately clear the infection. Experimental protocols often use lower concentrations, in which case the beneficial effects of collateral sensitivity could be substantial for rapid cycling regimens. Aulin et al. [21] indeed conclude that collateral sensitivity can be best exploited at rather low concentrations, which are lower than the ones that we consider.

Aulin et al. [21] find that collateral sensitivity reduces resistance evolution in the 1-day cycling regimen but has weak effects in the slower regimens, turning 1-day cycling superior to a single drug switch for drug pairs with a large Hill coefficient. Udekwu, and Weiss [20], on the other hand, find that slower cycling has an advantage over rapid cycling for drugs exhibiting reciprocal collateral sensitivity (they, unfortunately, do not compare to drugs without collateral sensitivity).

### The role of drug-drug interactions in treatment

Although drug-drug interactions are effective only in the overlap phases of doses from different drugs, our results show an influence of the type of interaction on the treatment efficiency and the relative performances of cycling frequencies. The time to suppress the bacterial population is shorter with a synergistic drug pair and longer with an antagonistic drug pair than in the absence of interactions as long as the collateral sensitivity profile is the same for both drug pairs. In that case, an antagonistic drug pair with collateral sensitivity effects can, however, sometimes be more efficient than a synergistic drug pair without such effects.

The influence of drug-drug interactions on treatment efficiency and resistance evolution has attracted much attention in the context of combination therapy, where both drugs are administered simultaneously. In line with our observations, synergistic interactions were found to increase bacterial clearance in *in vitro* combination treatment, whereas antagonism was associated with a low probability of population extinction [45]. On the other hand, other studies reported accelerated resistance evolution under synergism for certain antibiotic concentrations [49, 50]. Similarly, modelling studies demonstrate that in the presence of resource competition, antagonistic drug pairs can have a benefit over synergistic ones in terms of reduced resistance evolution, which is due to the strength of competitive release experienced by resistant types [50, 51]. With hyper-antagonism (suppressive drug-drug interactions), there might even be selection against resistant variants [52–55]. In a supplementary analysis (SI section S2.8), we found ranges of drug doses in which synergism leads to larger sizes of the resistant sub-populations than antagonism or Bliss independence. However, these doses were lower than the first effective dose and thus lower than the doses considered in the main text. Our observations for high doses accord with the finding of Rodriguez de Evgrafov et al. [19] who observed no influence of drug-drug interactions on resistance evolution under high antibiotic concentrations in a combination treatment in the lab.

### Limitations and extensions

In our comparison between the lab and the patient, we focused on the consequences of dose overlaps, which are present in the patient but not in the lab. However, this is, of course, not the only difference between the two environments. In the patient, the bacteria encounter a spatially structured environment, commensal bacteria, the human immune system etc..

On the side of the biology of bacteria, important factors that we did not include are hypermutator strains and stress-induced mutagenesis, which could lower the efficiency of sequential therapy. We further assumed that all bacteria in the population replicate at the same rate. However, different sub-populations might replicate at different rates and include persister types as, for example, observed by Kaiser et al. [56] in mice. We showed that the optimal strategy can vary with the replication rate of the bacteria, which could be of importance for the treatment of heterogeneous populations.

In our study, we focused on a limited set of drug pairs to understand the dynamics in some detail and to provide thorough explanations for the observed outcomes. We especially did not vary the pharmacokinetics of the drugs. In our example, the antibiotic has decayed to around 10% of the administered dose, when the next dose is given. If the intervals between administrations are short and/or the half lives of the drugs long, both drugs are simultaneously present in the body at similar concentrations during rapid cycling. This would resemble combination therapy with short intervals between administrations; combination therapy with long intervals between administrations is different in that drug levels might temporarily decrease to low levels. Comparing the two strategies combination and sequential therapy, both the modeling study by Aulin et al. [21] and the experimental study by Fuentes-Hernandez et al. [9] find conditions under which sequential therapy performs better than combination therapy and the other way round. This indicates that the comparison is not straightforward, and a detailed understanding would require a thorough analysis. We further only considered bactericidal antibiotics. For bacteriostatic drugs, the maximum kill rate is given by the intrinsic death rate. Moreover, since replication and mutation are coupled, a drug-induced decrease in the replication rate will also reduce the per-capita rate at which mutations occur. Consequently, the treatment efficiency and the optimal cycling regimen might depend on the mode of action of the drugs.

Previous work on sequential patient treatment is equally restricted to a few specific parameter sets [20, 21]. While this is a meaningful approach to gain first fundamental insights, future work is needed that provides parameter (and maybe also structural) sensitivity analyses to unite and synthesise results from different studies. A particular challenge in the comparison across different studies, which has also been highlighted in the context of drug cycling and other multi-drug therapies at a community level [4], is the use of different optimality criteria, since different criteria might identify different strategies as optimal.

## Conclusion

Our results show that sequential therapy efficiently controls the spread of resistance and suppresses the bacterial population. Optimal treatment settings depend on the replication rate of the bacteria. The optimal cycling frequency usually lies within the range of rapid cycling regimens. Switching the drug at every administration is, however, sometimes sub-optimal for patient treatment. At least when resistance does not pre-exist prior to treatment and antibiotic concentrations are well above the wild-type MIC, the benefits provided by collateral sensitivity are negligible for rapid cycling regimens. In contrast, drug-drug interactions, which have received little attention in the context of sequential therapy so far, substantially affect the treatment efficiency if drugs are quickly alternated, especially if the cell division rate is low. This suggests that drug-drug interactions could be an equally or even more important criterion than collateral sensitivity for the selection of drug pairs for clinical protocols of rapid cycling. On the other hand, when rapid cycling is not possible for logistic reasons or tolerability and slower regimens are applied, collateral sensitivity can be a better determinant of treatment efficiency than drug-drug interactions. Overall, we see that pharmacokinetic processes that occur in the patient but are absent in classic laboratory experiments cannot be ignored in the design of clinical treatment protocols.

## Supporting information

Additional information for the methods and results section of the main paper.

## Acknowledgements

We thank Ernesto Berríos-Caro for pointing us to Lewis’ thinning algorithm and for providing advice on the semi-stochastic simulations. We thank Roland Regös, Claudia Igler, Sebastian Bonhoeffer, Hinrich Schulenburg, and Aditi Batra, and the members of the Research group Stochastic Evolutionary Dynamics for helpful discussions.

## Data availability

The simulation code and the data is available at: https://gitlab.gwdg.de/nyhoegen/sequential-therapy-supplementary-information

## References

(1) Roemhild, R., and Schulenburg, H. (2019). Evolutionary ecology meets the antibiotic crisis: Can we control pathogen adaptation through sequential therapy? Evolution, Medicine, and Public Health 2019, 37–45.

(2) Andersson, D. I., Balaban, N. Q., Baquero, F., Courvalin, P., Glaser, P., Gophna, U., Kishony, R., Molin, S., and Tønjum, T. (2020). Antibiotic resistance: Turning evolutionary principles into clinical reality. FEMS Microbiology Reviews 44, 171–188.

(3) Merker, M., Tueffers, L., Vallier, M., Groth, E. E., Sonnenkalb, L., Unterweger, D., Baines, J. F., Niemann, S., and Schulenburg, H. (2020). Evolutionary approaches to combat antibiotic resistance: opportunities and challenges for precision medicine. Frontiers in Immunology 11, 1938.

(4) Uecker, H., and Bonhoeffer, S. (2021). Antibiotic treatment protocols revisited: the challenges of a conclusive assessment by mathematical modelling. Journal of The Royal Society Interface 18, 20210308.

(5) Kim, S., Lieberman, T. D., and Kishony, R. (2014). Alternating antibiotic treatments constrain evolutionary paths to multidrug resistance. Proceedings of the National Academy of Sciences 111, 14494–14499.

(6) Roemhild, R., Barbosa, C., Beardmore, R. E., Jansen, G., and Schulenburg, H. (2015). Temporal variation in antibiotic environments slows down resistance evolution in pathogenic *Pseudomonas aeruginosa*. Evolutionary Applications 8, 945–955.

(7) Yoshida, M., Reyes, S. G., Tsuda, S., Horinouchi, T., Furusawa, C., and Cronin, L. (2017). Time-programmable drug dosing allows the manipulation, suppression and reversal of antibiotic drug resistance in vitro. Nature Communications 8, 1–11.

(8) Batra, A., Roemhild, R., Rousseau, E., Franzenburg, S., Niemann, S., and Schulenburg, H. (2021). High potency of sequential therapy with only *β*-lactam antibiotics. eLife 10, e68876.

(9) Fuentes-Hernandez, A., Plucain, J., Gori, F., Peña-Miller, R., Reding, C., Jansen, G., Schulenburg, H., Gudelj, I., and Beardmore, R. E. (2015). Using a sequential regimen to eliminate bacteria at sublethal antibiotic dosages. PLoS Biology 13, e1002104.

(10) Roemhild, R., Gokhale, C. S., Dirksen, P., Blake, C., Rosenstiel, P., Traulsen, A., Andersson, D. I., and Schulenburg, H. (2018). Cellular hysteresis as a principle to maximize the efficacy of antibiotic therapy. Proceedings of the National Academy of Sciences 115, 9767–9772.

(11) Imamovic, L., and Sommer, M. O. A. (2013). Use of collateral sensitivity networks to design drug cycling protocols that avoid resistance development. Science Translational Medicine 5, 204ra132–204ra132.

(12) Maltas, J., and Wood, K. B. (2019). Pervasive and diverse collateral sensitivity profiles inform optimal strategies to limit antibiotic resistance. PLoS Biology 17, e3000515.

(13) Szybalski, W., and Bryson, V. (1952). Genetic studies on microbial cross resistance to toxic agents I. Cross resistance of *Escherichia coli* to fifteen antibiotics. Journal of Bacteriology 64, 489–499.

(14) Lázár, V., Pal Singh, G., Spohn, R., Nagy, I., Horváth, B., Hrtyan, M., Busa-Fekete, R., Bogos, B., Méhi, O., Csörgő, B., et al. (2013). Bacterial evolution of antibiotic hypersensitivity. Molecular Systems Biology 9, 700.

(15) Podnecky, N. L., Fredheim, E. G. A., Kloos, J., Sørum, V., Primicerio, R., Roberts, A. P., Rozen, D. E., Samuelsen, Ø., and Johnsen, P. J. (2018). Conserved collateral antibiotic susceptibility networks in diverse clinical strains of *Escherichia coli*. Nature Communications 9, 1–11.

(16) Munck, C., Gumpert, H. K., Nilsson Wallin, A. I., Wang, H. H., and Sommer, M. O. A. (2014). Prediction of resistance development against drug combinations by collateral responses to component drugs. Science Translational Medicine 6, 262ra156–262ra156.

(17) Barbosa, C., Trebosc, V., Kemmer, C., Rosenstiel, P., Beardmore, R. E., Schulenburg, H., and Jansen, G. (2017). Alternative evolutionary paths to bacterial antibiotic resistance cause distinct collateral effects. Molecular Biology and Evolution 34, 2229–2244.

(18) Yen, P., and Papin, J. A. (2017). History of antibiotic adaptation influences microbial evolutionary dynamics during subsequent treatment. PLoS Biology 15, e2001586.

(19) Rodriguez de Evgrafov, M., Gumpert, H., Munck, C., Thomsen, T. T., and Sommer, M. O. A. (2015). Collateral resistance and sensitivity modulate evolution of high-level resistance to drug combination treatment in *Staphylococcus aureus*. Molecular Biology and Evolution 32, 1175–1185.

(20) Udekwu, K. I., and Weiss, H. (2018). Pharmacodynamic considerations of collateral sensitivity in design of antibiotic treatment regimen. Drug Design, Development and Therapy 12, 2249.

(21) Aulin, L., Liakopoulos, A., van der Graaf, P. H., Rozen, D. E., and van Hasselt, J. G. (2021). Design principles of collateral sensitivity-based dosing strategies. Nature Communications 12, 1–14.

(22) Regoes, R. R., Wiuff, C., Zappala, R. M., Garner, K. N., Baquero, F., and Levin, B. R. (2004). Pharmacodynamic functions: a multiparameter approach to the design of antibiotic treatment regimens. Antimicrobial Agents and Chemotherapy 48, 3670–3676.

(23) Chevereau, G., Dravecká, M., Batur, T., Guvenek, A., Ayhan, D. H., Toprak, E., and Bollenbach, T. (2015). Quantifying the determinants of evolutionary dynamics leading to drug resistance. PLoS Biology 13, e1002299.

(24) Das, S. G., Direito, S. O. L., Waclaw, B., Allen, R. J., and Krug, J. (2020). Predictable properties of fitness landscapes induced by adaptational tradeoffs. eLife 9, e55155.

(25) Wicha, S. G., Chen, C., Clewe, O., and Simonsson, U. S. H. (2017). A general pharma-codynamic interaction model identifies perpetrators and victims in drug interactions. Nature Communications 8, 1–11.

(26) Roemhild, R., Bollenbach, T., and Andersson, D. I. (2022). The physiology and genetics of bacterial responses to antibiotic combinations. Nature Reviews Microbiology 20, 478–490.

(27) Bliss, C. I. (1939). The toxicity of poisons applied jointly 1. Annals of Applied Biology 26, 585–615.

(28) Baeder, D. Y., Yu, G., Hoze, N., Rolff, J., and Regoes, R. R. (2016). Antimicrobial combinations: Bliss independence and Loewe additivity derived from mechanistic multi-hit models. Philosophical Transactions of the Royal Society B: Biological Sciences 371, 20150294.

(29) Levison, M. E., and Levison, J. H. (2009). Pharmacokinetics and pharmacodynamics of antibacterial agents. Infectious Disease Clinics 23, 791–815.

(30) Shargel, L., Wu-Pong, S., and Yu, A. B. C., Applied biopharmaceutics & pharmacokinetics, 5th ed.; McGraw-Hill Professional Publishing: New York, 2007.

(31) Shampine, L. F., and Reichelt, M. W. (1997). The matlab ODE suite. SIAM Journal on Scientific Computing 18, 1–22.

(32) Lewis, P. A. W., and Shedler, G. S. (1979). Simulation of nonhomogeneous Poisson processes by thinning. Naval Research Logistics Quarterly 26, 403–413.

(33) Gillespie, D. T. (1976). A general method for numerically simulating the stochastic time evolution of coupled chemical reactions. Journal of Computational Physics 22, 403–434.

(34) Gillespie, D. T. (1977). Exact stochastic simulation of coupled chemical reactions. The Journal of Physical Chemistry 81, 2340–2361.

(35) Kiehl, T. R., Mattheyses, R. M., and Simmons, M. K. (2004). Hybrid simulation of cellular behavior. Bioinformatics 20, 316–322.

(36) Rodríguez-Rojas, A., Makarova, O., and Rolff, J. (2014). Antimicrobials, stress and mutagenesis. PLoS Pathogens 10, e1004445.

(37) Imhof, M., and Schlötterer, C. (2001). Fitness effects of advantageous mutations in evolving *Escherichia coli* populations. Proceedings of the National Academy of Sciences 98, 1113–1117.

(38) Grant, A. J., Restif, O., McKinley, T. J., Sheppard, M., Maskell, D. J., and Mastroeni, P. (2008). Modelling within-host spatiotemporal dynamics of invasive bacterial disease. PLoS Biology 6, e74.

(39) Igler, C., Rolff, J., and Regoes, R. (2021). Multi-step vs. single-step resistance evolution under different drugs, pharmacokinetics, and treatment regimens. eLife 10, e64116.

(40) Spohn, R., Daruka, L., Lázár, V., Martins, A., Vidovics, F., Grézal, G., Méhi, O., Kintses, B., Számel, M., Jangir, P. K., et al. (2019). Integrated evolutionary analysis reveals antimicrobial peptides with limited resistance. Nature Communications 10, 1–13.

(41) Melnyk, A. H., Wong, A., and Kassen, R. (2015). The fitness costs of antibiotic resistance mutations. Evolutionary Applications 8, 273–283.

(42) Vance-Bryan, K., Guay, D. R. P., and Rotschafer, J. C. (1990). Clinical pharmacokinetics of ciprofloxacin. Clinical Pharmacokinetics 19, 434–461.

(43) Naber, K. G., Theuretzbacher, U., Moneva-Koucheva, G., and Stass, H. (1999). Urinary excretion and bactericidal activity of intravenous ciprofloxacin compared with oral ciprofloxacin. European Journal of Clinical Microbiology and Infectious Diseases 18, 783–789.

(44) Kaiser, P., Slack, E., Grant, A. J., Hardt, W.-D., and Regoes, R. R. (2013). Lymph node colonization dynamics after oral *Salmonella* Typhimurium infection in mice. PLoS Pathogens 9, e1003532.

(45) Barbosa, C., Beardmore, R. E., Schulenburg, H., and Jansen, G. (2018). Antibiotic combination efficacy (ACE) networks for a *Pseudomonas aeruginosa* model. PLoS Biology 16, e2004356.

(46) Gjini, E., and Wood, K. B. (2021). Price equation captures the role of drug interactions and collateral effects in the evolution of multidrug resistance. Elife 10, e64851.

(47) Greenfield, B. K., Shaked, S., Marrs, C. F., Nelson, P., Raxter, I., Xi, C., McKone, T. E., and Jolliet, O. (2018). Modeling the emergence of antibiotic resistance in the environment: an analytical solution for the minimum selection concentration. Antimicrobial Agents and Chemotherapy 62, e01686–17.

(48) Yu, G., Baeder, D. Y., Regoes, R. R., and Rolff, J. (2018). Predicting drug resistance evolution: insights from antimicrobial peptides and antibiotics. Proceedings of the Royal Society B: Biological Sciences 285, 20172687.

(49) Hegreness, M., Shoresh, N., Damian, D., Hartl, D., and Kishony, R. (2008). Accelerated evolution of resistance in multidrug environments. Proceedings of the National Academy of Sciences 105, 13977–13981.

(50) Peña-Miller, R., Laehnemann, D., Jansen, G., Fuentes-Hernandez, A., Rosenstiel, P., Schulenburg, H., and Beardmore, R. E. (2013). When the most potent combination of antibiotics selects for the greatest bacterial load: the smile-frown transition. PLoS Biology 11, e1001540.

(51) Torella, J. P., Chait, R., and Kishony, R. (2010). Optimal drug synergy in antimicrobial treatments. PLoS Computational Biology 6, e1000796.

(52) Yeh, P. J., Hegreness, M. J., Presser Aiden, A., and Kishony, R. (2009). Drug interactions and the evolution of antibiotic resistance. Nature Reviews Microbiology 7, 460–466.

(53) Bollenbach, T. (2015). Antimicrobial interactions: mechanisms and implications for drug discovery and resistance evolution. Current Opinion in Microbiology 27, 1–9.

(54) Baym, M., Stone, L. K., and Kishony, R. (2016). Multidrug evolutionary strategies to reverse antibiotic resistance. Science 351.

(55) Singh, N., and Yeh, P. (2017). Suppressive drug combinations and their potential to combat antibiotic resistance. The Journal of Antibiotics 70, 1033–1042.

(56) Kaiser, P., Regoes, R. R., Dolowschiak, T., Wotzka, S. Y., Lengefeld, J., Slack, E., Grant, A. J., Ackermann, M., and Hardt, W.-D. (2014). Cecum lymph node dendritic cells harbor slow-growing bacteria phenotypically tolerant to antibiotic treatment. PLoS Biology 12, e1001793.

