## Additional information for the methods and results section of the main paper. for "Sequential antibiotic therapy in the lab and in the patient"

### S1 Supporting information on the methods

#### S1.1 Including drug-drug interactions into the pharmacodynamic function

To account for drug-drug interactions, we included the generalized pharmacodynamic interaction term introduced by Wicha et al. [1] into the PD-model by Regoes et al. [2], which we briefly outline here.

Both studies have the $E_{\max}$ model as a starting point. In this model, the effect of a drug is described by the term

$$E(c) = \frac{E_{\max} \cdot c^{\kappa}}{\mathrm{EC}_{50}^{\kappa} + c^{\kappa}}, \tag{S1}$$

where $E_{\max}$ is the maximal possible effect and $\mathrm{EC}_{50}$ the drug concentration that produces 50% of the maximum effect. $\kappa$ is the Hill coefficient. (For our study, the effect of the drug is the antibiotic-induced kill rate, i.e. $E(c) = \mu(c)$.)

The drug interaction term $I(c)$ introduced by Wicha et al. [1] describes how a second drug alters the effect of a first drug and is defined by

$$I(c') = \frac{\mathrm{INT} \cdot c'^{H}}{\mathrm{EC}_{50,\mathrm{Int}}^{H} + c'^{H}}, \tag{S2}$$

where $c'$ is the concentration of the second drug, $\mathrm{INT}$ is the maximal effect that it can have, $\mathrm{EC}_{50,\mathrm{Int}}$ is the concentration to reach $50\%$ of $\mathrm{INT}$, and $H$ is the steepness of the curve. Depending on the drugs, the interaction term can either alter $E_{\max}$ or $\mathrm{EC}_{50}$. In our study, it alters $\mathrm{EC}_{50}$.

The effect of a drug A, influenced by the drug-drug-interaction caused by a drug B, is calculated by substituting $EC_{50}$ with the term $\mathrm{EC}_{50} \cdot (1 + I(c_B))$:

$$E(c_A) = \frac{E_{\max,A} \cdot c_A^{\kappa_A}}{\mathrm{EC}_{50,A}^{\kappa_A} \cdot \left(1 + \frac{\mathrm{INT} \cdot c_B^H}{\mathrm{EC}_{50,\mathrm{Int}}^H + c_B^H}\right)^{\kappa_A} + c_A^{\kappa_A}}, \tag{S3}$$

where we gave a subscript $A$ to the parameters of the $E_{\max}$ model.

The parameter $\mathrm{INT}$ can range from -1 to infinity. For a value smaller than zero, drug B potentiates the effect of drug A; for a value larger than zero, drug B inhibits drug A. If $\mathrm{INT} = 0$, drug B does not alter the effect of drug A.

The PD-model by Regoes et al. [2] – Eq. (2) in our manuscript – is derived from the general $E_{\max}$ model; the two models can be translated into each other with two substitutions:

- $E_{\max} = \psi_{\max} - \psi_{\min}$
- $\mathrm{EC}_{50}^{\kappa} = \frac{-\psi_{\min} \cdot \mathrm{zMIC}^{\kappa}}{\psi_{\max}}$

Multiplication of $\mathrm{EC}_{50}$ with the term $(1+I(c'))$ thus leads to a multiplication of the parameter $\mathrm{zMIC}$ with $(1 + I(c'))$.

#### S1.2 The pharmacodynamic surface under antagonistic drug-drug interactions

Much experimental work has been invested into quantifying, classifying, and understanding drug-drug interactions across concentration gradients [e.g. 3, 4, 5, 6, 7]. On the theoretical side, several approaches have been developed to describe the pharmacodynamics of drug-drug interactions [4, 8, 1, 9]. None of them accurately captures all observed fitness landscapes. For our 'proof-of-principle' study, it is sufficient to decide on one implementation of drug-drug interaction as an example. More detailed future models could explore a larger range of implementations or be based on empirically measured pharmacodynamic surfaces.

Figure S1 illustrates some features of our implementation of drug-drug interactions (Eq. (5)). Panels A and E compare the antibiotic-induced kill rate of the wild type in the presence and absence of drug B. We see that there is a considerable range in which the antibiotic-induced kill rate is larger when only drug A is present than in the presence of both drugs. The other panels compare the effect of the combined drugs on the mutant types with the effect on the susceptible wild type. These panels show that the wild type is always more affected by the treatment than the mutants.

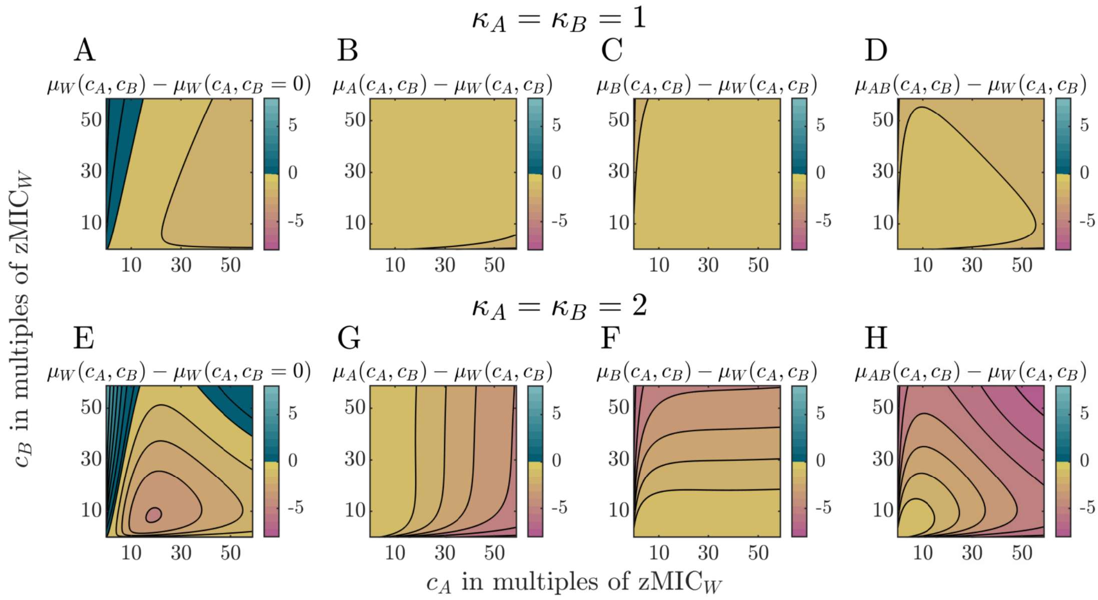


Figure S1: Comparison of the pharmacodynamic surfaces for different cell types under antagonistic drug-drug interactions. Panels A ($\kappa_A = \kappa_B = 1$) and E ($\kappa_A = \kappa_B = 2$) show for the wild type the difference in the antibiotic-induced kill rate between the combination treatment and mono-therapy with drug A. The comparison shows regions for which the mono-therapy leads to higher kill rates than the combination (yellow and pink colors). Panels B-C ($\kappa_A = \kappa_B = 1$) and F-H ($\kappa_A = \kappa_B = 2$) compare the antibiotic-induced kill rates of the mutant types with that of the wild type. Each comparison shows a higher kill rate for the wild type (yellow and pink colors). The parameter values were chosen according to Table 1.

#### S1.3 Description of the semi-stochastic hybrid model

In this section, we explain the implementation of our stochastic-deterministic hybrid model in more detail. The algorithm is additionally displayed in pseudo-code below (see page 5). The implementation is done in MATLAB (version 2020a/2020b).

The stochastic part of our hybrid model is simulated with the Lewis' Thinning algorithm (LTA) [10], which can be used as an extension of the Gillespie algorithm [11, 12] when rates are time-dependent. The algorithm simulates the population growth one event – i.e. cell replication (with or without mutation) or death – at a time. The waiting time $\tau$ until the next event occurs is drawn from an exponential distribution, where the parameter of the distribution depends on the cell numbers and an upper bound for the time-dependent rates at which the events occur. We use a new estimate for the upper bound at every time step. For the replication rates (with or without mutation), we choose the intrinsic replication rate $r_0$ as an upper bound. For the kill rates, we use the value at the current time point. The only exception we need to make is when the current time point is very close to an antibiotic administration. Whenever the current time point is within 0.9 hours before the next administration, we use the kill rates at dose $D$ as upper bound. By trial, the value of 0.9 hours was found to be large enough that the jump in the kill rate at the administration is not missed and small enough that we do not choose an unnecessarily high upper bound too often. Based on the actual rates at the new time point (after step $\tau$), denoted by $s$ in the pseudo-code below, it is decided whether an event occurs. In case of acceptance, based on the respective rates, the type of event (replication with or without mutation, death) is chosen with the respective probability. The cell numbers and the time $t$ are updated accordingly.

To increase the speed of the algorithm, we are following the hybrid approach described in Kiehl et al. [13] and only simulate the replication or death events for types with a population size below a certain threshold ($N_{\text{det}}$, here set to 1000) and mutations stochastically. Whenever the size of a sub-population exceeds the threshold, we determine the cell numbers for a given time point by solving the deterministic ODE system for that type. A state vector ($S$) indicates for each type with a one (or a zero) whether the respective sub-population size is above (or below) the threshold $N_{\text{det}}$. For each possible state vector, we define a set of possible events for the stochastic part and a system of ordinary differential equations (ODE-system) for the deterministic part, which only includes the types with sub-population sizes above the threshold. When all sub-populations drop below the threshold ($S = [0, 0, 0, 0]$), everything is stochastic. For the other cases, the update in cell numbers goes as follows: first, the time step $\tau$ is calculated based on the subset of possible events (depending on the state $S$). It is possible that $\tau$ is quite large, and the next stochastic event will only occur after a long time. We still want to keep track of the deterministic populations in those cases. Therefore, we define a deterministic time-step ($\tau_{\text{det}} = 1$), which is used to update the deterministic sub-populations, whenever $\tau$ exceeds $\tau_{\text{det}}$. The sub-populations are updated by solving the ODE system for the given time interval. The total cell number, the time $t$, and the system's state are updated accordingly. If the time step $\tau$ is smaller than the deterministic time step $\tau_{\text{det}}$, the deterministic sub-populations are updated first. Then, following the LTA, it is decided if the time step gets accepted. When it gets accepted, the cell numbers, the vector $S$ and the time variable $t$ will be updated (the time variable $s$ is always updated by $\tau$). The algorithm terminates when a certain criterion is met. Here, we terminate the algorithm when the maximum time or a population size of zero is reached.

Algorithm 1: Pseudocode for the implementation of the Hybrid-model.

Initialize parameter:
$W = N_0$, $M_A = 0$, $M_B = 0$, $M_{AB} = 0$, $N_{\text{all}} = N_0$
$t = 0$, $s = 0$ ▷ initialize two variables counting the time
$S = [1, 0, 0, 0]$ ▷ set states according to the subpopulation sizes
$\tau_{\text{det}} = 1$ ▷ set size of discrete time steps
$N_{\text{det}} = 1000$ ▷ set threshold for stochastic simulations

When at least one subpopulation size exceeds $N_{\text{det}}$:
**while** $s < t_{\text{max}}$ & $N_{\text{all}} > 0$ & $S \neq [0, 0, 0, 0]$ **do**
  Calculate time step $\tau$ according to LTA
  **if** $\tau > \tau_{\text{det}}$ **then**
    Set $s = s + \tau_{\text{det}}$
    Update only the deterministic sub-populations:
    Solve respective ODE-system for the interval $[s - \tau_{\text{det}}, s]$
    Update $N_{\text{all}}$ and set $t = t + \tau_{\text{det}}$
    Set $S = ([W, M_A, M_B, M_{AB}] > N_{\text{det}})$
    Save all cell numbers for the time point $t$
  **else**
    Set $s = s + \tau$
    Update first the deterministic sub-populations:
    Solve respective ODE-system for the interval $[s - \tau, s]$
    Decide with LTA if an event occurs
    **if** event occurs **then**
      Decide with LTA which event occurs
      Update respective type, $N_{\text{all}}$ and set $t = t + \tau$
      Save all cell numbers for the time point $t$
    **end if**
    Set $S = ([W, M_A, M_B, M_{AB}] > N_{\text{det}})$
  **end if**
**end while**

When all subpopulation sizes are below $N_{\text{det}}$:
**while** $s < t_{\text{max}}$ & $N_{\text{all}} > 0$ & $S = [0, 0, 0, 0]$ **do**
  Update cell numbers and time according to LTA
  Set $S = ([W, M_A, M_B, M_{AB}] > N_{\text{det}})$
  Save all cell numbers for the time point $t$
**end while**

#### S1.4 Comparison of results obtained from the deterministic model and the semi-stochastic hybrid model

We want to make sure that conclusions derived from the deterministic ODE model are robust with respect to stochasticity in cell birth and death and mutation events. To test the robustness of predictions from the ODE model, we performed simulations using the semi-stochastic hybrid model for a small set of treatments and compared the time until all bacterial sub-population sizes have dropped to a threshold size $N_c$ (our primary measure) with the outcome of the deterministic model (Figure S2).

Overall, the results of the stochastic and the deterministic model align well. However, with increasing concentrations, the agreement gets less good. Especially, the deterministic model suggests an abrupt change in treatment efficiency at some concentration (best visible in panels H-J), whereas the treatment efficiency changes smoothly in the stochastic model.

The reason for this sudden drop in the time until all sub-population sizes are suppressed below a threshold $N_c$ (our measure of treatment efficiency) is illustrated in Figure S3, which shows the cell dynamics slightly to the left (Panel A) and slightly to the right (Panel B) of the drop. The wild-type cell numbers are decreasing, while the cell numbers of the type that is resistant to drug $A$ are increasing. In Panel B, the wild-type population size falls below the threshold $N_c$ before the sub-population size of the single resistant mutant is above $N_c$. In Panel A, the antibiotic dose is slightly lower such that the wild-type population size declines a bit slower, and the single mutants grow a bit faster. As a consequence, the wild-type population size fails to fall below the threshold before the single mutant grows above $N_c$, and single mutants grow until the drug is switched to drug B. In Figure 3 in the main text, to the right of the drop, two administrations of drug A are sufficient to suppress all sub-population sizes below $N_c$ (having a situation as in Panel A of Figure S3). In that case, all cycling regimens except for the fastest one, in which drugs are switched at every administration, are equivalent.

In the semi-stochastic simulation, no such drop in the time for all sub-populations to reach $N_c$ occurs since the occurrence of mutations (and their early dynamics) are stochastic. They do not occur (and spread) with certainty below a given concentration while being impossible above it. They rather occur (and spread) by chance with a probability that changes gradually across the dose gradient.

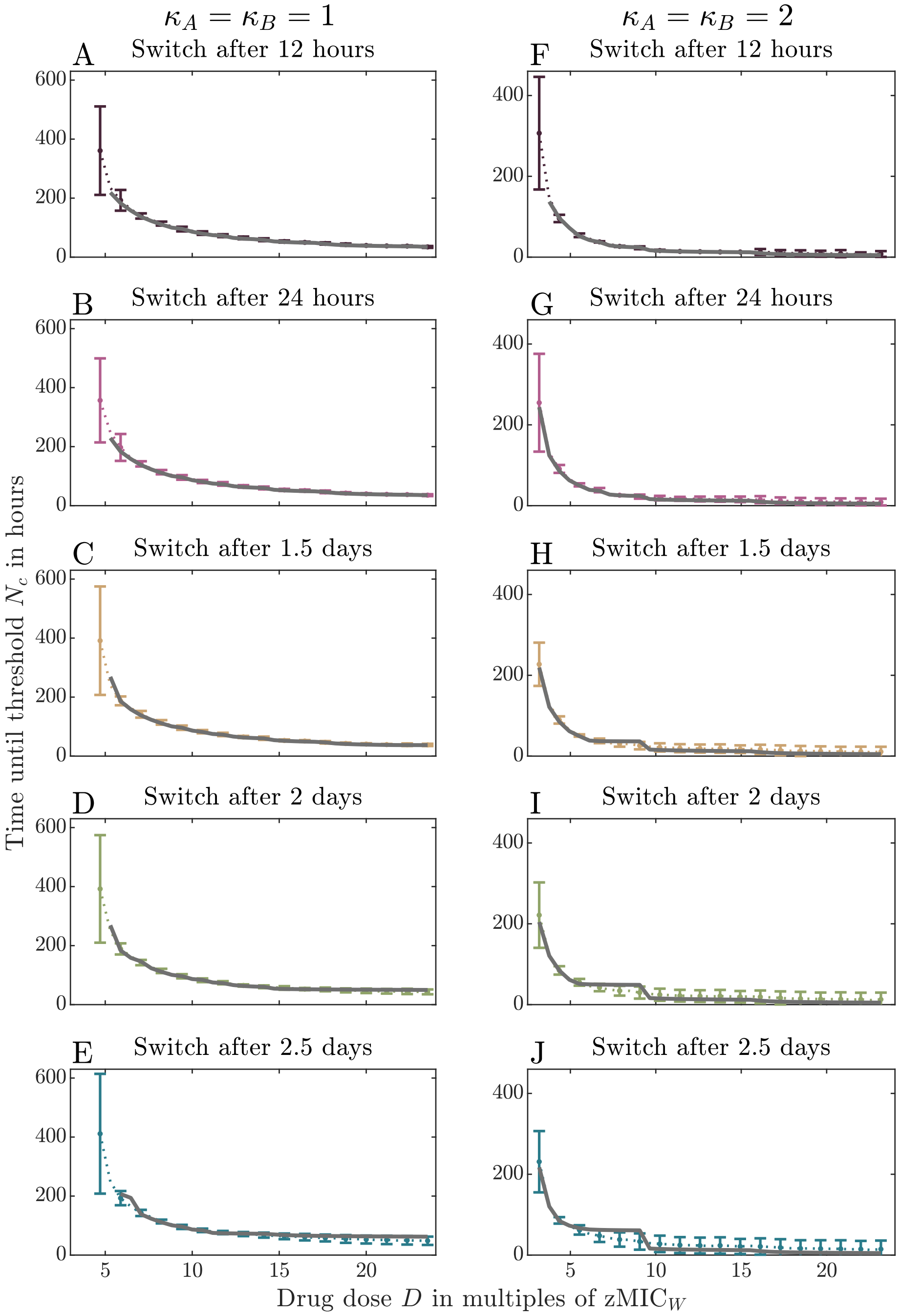


Figure S2: Comparison of results from the deterministic ODE model and the semi-stochastic hybrid model. Each plot displays the time until all bacterial sub-population sizes have dropped to the threshold size $N_c$ as a function of the drug dose $D$ at administration for a specific treatment regimen (indicated by the title). In the left column (A-E), the drugs have Hill coefficients $\kappa_A = \kappa_B = 1$; in the right column (F-J), the drugs have Hill coefficients $\kappa_A = \kappa_B = 2$. Note the different scales of the $y$-axis in the two columns. The grey line shows the solution from the deterministic ODE model; the colored symbols show the mean of 100 replicate runs of the semi-stochastic hybrid model. Error bars indicate the standard deviation (not the standard error) to demonstrate the variability between replicates.

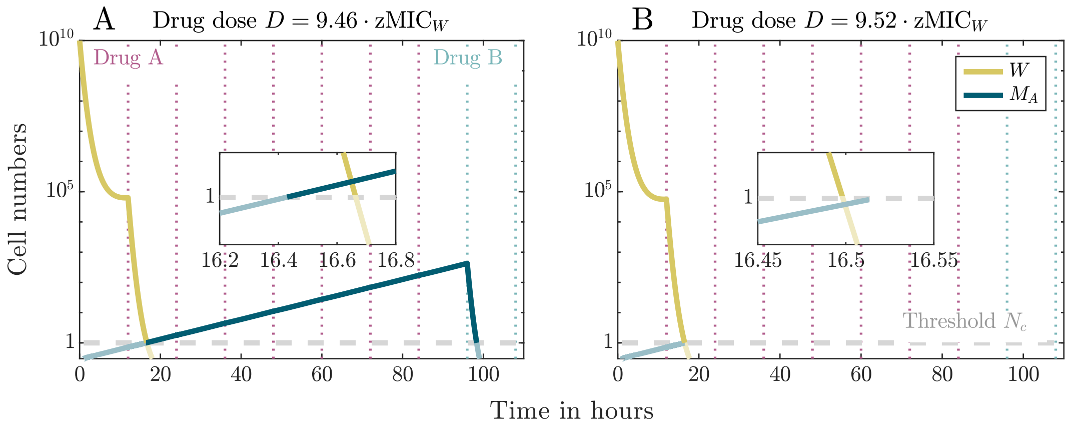


Figure S3: Cell dynamics for concentrations slightly to the left and to the right of the sudden change in treatment efficiency for $\kappa_A = \kappa_B = 2$. The time between the drug switches is four days. Slightly to the left of the sudden change (Panel A), it is necessary to wait for a switch to drug B to control the single resistant mutant. Slightly to the right (Panel B), the bacterial population declines a little bit more quickly and suppression is achieved within two drug administrations.

#### S1.5 The variation across patients

With the semi-stochastic model, we can assess not only the mean efficiency of a treatment but also the variation across patients. For high doses, the standard deviation in the time for all sub-populations to reach $N_c$ increases with decreasing cycling frequency (the size of the error bars increases from top to bottom in Figure S2). This implies that the fast cycling regimens are not only better than slow cycling regimens on average but also more reliable as they are less affected by stochasticity for high concentrations. For low doses, the standard deviation is large for all treatments. Figure S4 shows that the dependency on the antibiotic dose and the cycling frequency is complex.

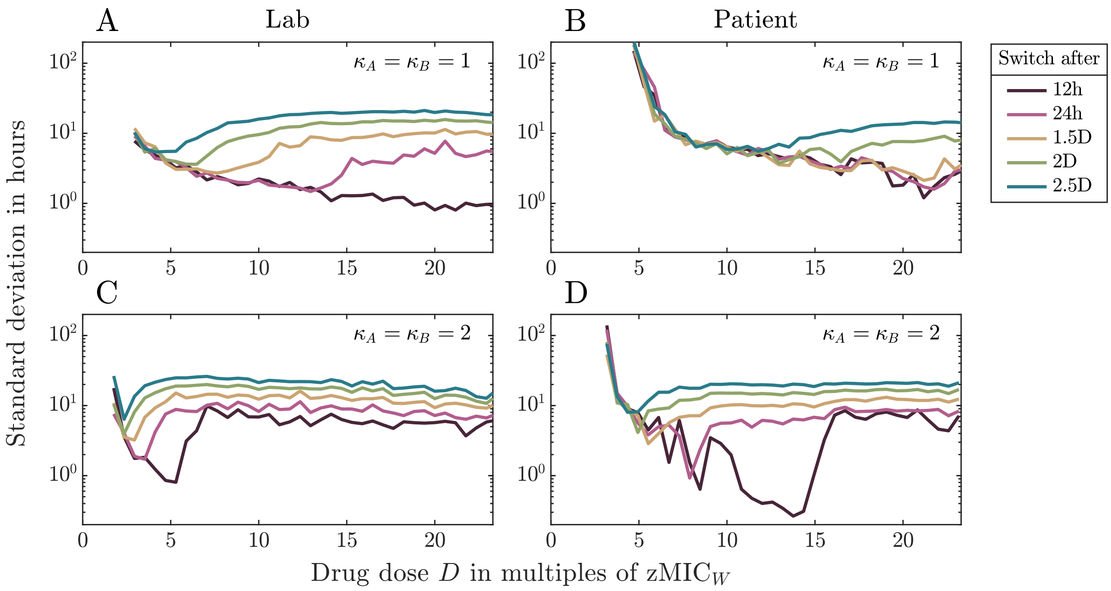


Figure S4: Standard deviation for simulations of five different cycling treatments across a wide range of concentrations. The left column shows the standard deviation of simulations in the lab environment and the right column in the patient environment. Panels A and B used drugs with $\kappa_A = \kappa_B = 1$ and panels C and D with $\kappa_A = \kappa_B = 2$. For each concentration, we ran 100 replicate simulations. Data points are connected by lines to guide the eye.

#### S2 Supporting information on the results for the treatment of slowly replicating bacteria

##### S2.1 Effect of $\psi_{\min}$ on treatment

Figure S5A displays the growth rate of the susceptible type for two drugs with different $\psi_{\min}$. The lines only start to deviate from each other for high concentrations, which exceed the concentration range that we considered in our analysis. As the two pharmacodynamic functions result in more or less the same growth rate for the considered concentrations, it is not surprising that there is no visible difference in the first effective concentration for two drug pairs that only differ in $\psi_{\min}$ (Panel B). For larger differences between the $\psi_{\min}$-values than considered in Figure S5, visible differences would likely appear.
Udekwu, and Weiss [14] found an advantage of drugs with lower $\psi_{\min}$ in a sequential regimen with collateral sensitivity. This indicates that $\psi_{\min}$ might play a role in other parts of the parameter space.

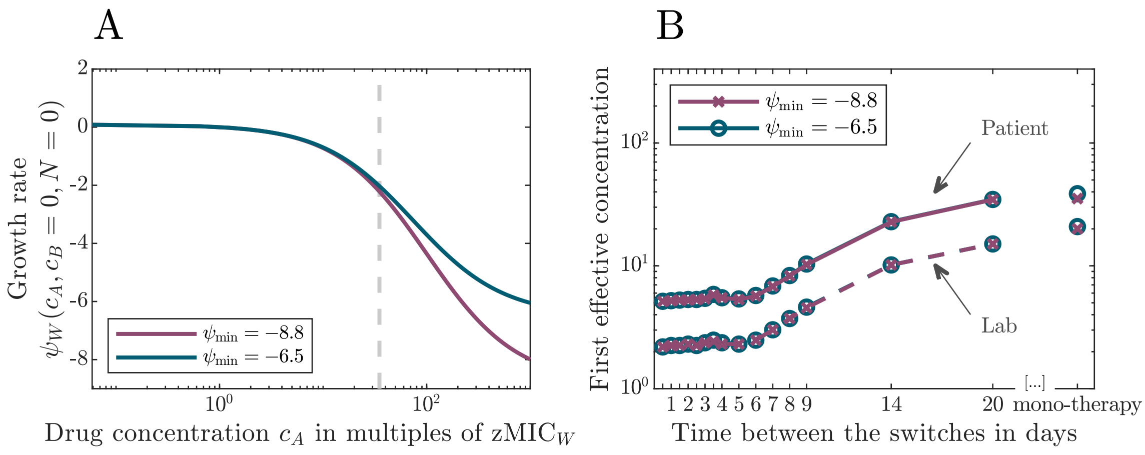


Figure S5: The effect of the minimal growth rate $\psi_{\min}$ on treatment. Panel A shows the pharmacodynamic function for two different values of $\psi_{\min}$ for $\kappa = 1$. The vertical grey line indicates the maximum concentration that we consider for treatment. Panel B compares the first effective concentrations of sequential therapy for the two values of $\psi_{\min}$ in the lab and in the patient as a function of the time between drug switches.

#### S2.2 Treatment with drugs with different pharmacodynamic parameters

In the main text, we assumed for simplicity that drug A and drug B have the same Hill coefficient, i.e. $\kappa_A = \kappa_B$. Here, we consider treatment with a mixed drug set with $\kappa_A \neq \kappa_B$ and compare it to treatment with drug pairs with equal Hill coefficients. The mixed drug sets differ by the order in which the drugs are applied, i.e. by which drug is applied first. Figure S6 shows the time until the threshold $N_c$ is reached by all sub-populations for the various drug pairs. The efficiency of the treatments with mixed drug pairs lies between the efficiencies for drug pairs with the same Hill coefficients. When the drugs are switched at every administration (Panel A), the treatments perform for high concentrations similar to the cycling of a drug pair with $\kappa_A = \kappa_B = 2$, irrespective of the starting drug. For intermediate and slow regimens (panels B and C), the efficiency at high concentrations is determined by the drug of the first administration.

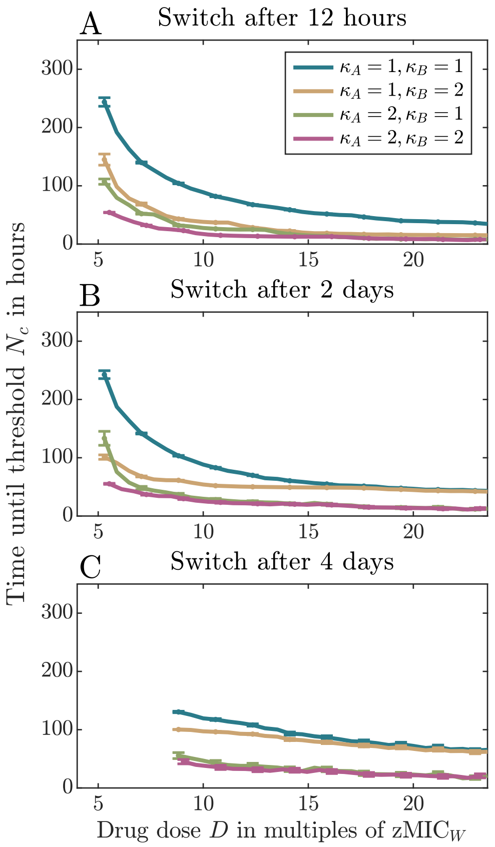


Figure S6: Efficiency of treatment with different drug pairs for three cycling regimens. Each Plot shows the time until the threshold $N_c$ is reached by all sub-populations as a function of the drug dose at administration. The Hill coefficients of the drugs can be the same for both drugs (blue and lilac) or differ between the drugs (orange and green). The treatment efficiency with mixed drug pairs is intermediate between those of drug pairs with equal Hill coefficients. For high concentrations, it converges to the efficiency of one of the 'pure' drug pairs. For very rapid cycling (Panel A), the efficiency in the limit of high concentrations is determined by the drug with the higher Hill coefficient, irrespective of the order of drugs in the mixed pair; for intermediate or slow cycling (panels B and C), the Hill coefficient of the drug at the first administration is critical.

#### S2.3 Antibiotic-induced killing during periods of dose overlap

In the main text, we saw that for low concentrations, there is a slight advantage of cycling the drugs less often in patient treatment when the drugs have a 'large' Hill coefficient (concretely, $\kappa_A = \kappa_B = 2$ in our example), while for a 'low' Hill coefficient ($\kappa_A = \kappa_B = 1$ in our example), the most rapid cycling is the optimal strategy.

We here show that this is due to the effects of dose overlaps from the same or from different drugs. To understand how such dose overlaps may influence the efficiency of the different treatments, we consider the antibiotic-induced kill rates in the presence of one drug or two drugs. For $\kappa_A = \kappa_B = 1$, the kill rate of the susceptible wild type is higher if drug A and drug B are present at a concentration $c_A = c_B = c$ each than if drug A is present at concentration $c_A = 2c$ and $c_B = 0$ or the other way round (Figure S7A). For $\kappa_A = \kappa_B = 2$, however, there exists a threshold concentration below which adding two doses of the same drug leads to a higher kill rate than giving two different drugs (Figure S7A). More generally, it can be shown mathematically that (1) for $\kappa_A = \kappa_B = \kappa \leq 1$, giving both drugs at concentration $c$ each always leads to a higher kill rate giving a single drug at concentration $2c$ and (2) for $\kappa_A = \kappa_B = \kappa > 1$, there exists a concentration below which giving two different drugs at concentration $c$ each leads to a lower kill rate than giving one drug at concentration $2c$, while it is the other way round above this threshold concentration.

For this, we compare the antibiotic induced kill rates $\mu(2c)$ (giving one drug at concentration $2c$) and $2\mu(c)$ (giving both drugs at concentration $c$ each), where the kill rate is given by Eq. (2), and ask when the ratio $\frac{\mu(2c)}{2\mu(c)}$ is larger than $1$:

$$\frac{\mu(2c)}{2\mu(c)} = \frac{\left(\frac{2c}{\text{zMIC}}\right)^{\kappa} \cdot \left\{\left(\frac{c}{\text{zMIC}}\right)^{\kappa} - \frac{\psi_{\min}}{\psi_{\max}}\right\}}{\left\{\left(\frac{2c}{\text{zMIC}}\right)^{\kappa} - \frac{\psi_{\min}}{\psi_{\max}}\right\} \cdot 2\left(\frac{c}{\text{zMIC}}\right)^{\kappa}} = \frac{1}{2}\,\frac{\left(\frac{2c}{\text{zMIC}}\right)^{\kappa} - 2^{\kappa}\frac{\psi_{\min}}{\psi_{\max}}}{\left(\frac{2c}{\text{zMIC}}\right)^{\kappa} - \frac{\psi_{\min}}{\psi_{\max}}} \overset{!}{>} 1 \tag{S4a}$$

$$\Leftrightarrow \left(\frac{2c}{\text{zMIC}}\right)^{\kappa} - 2^{\kappa}\frac{\psi_{\min}}{\psi_{\max}} \overset{!}{>} 2\left(\frac{2c}{\text{zMIC}}\right)^{\kappa} - 2\frac{\psi_{\min}}{\psi_{\max}} \tag{S4b}$$

$$\Leftrightarrow (2 - 2^{\kappa}) \cdot \frac{\psi_{\min}}{\psi_{\max}} \overset{!}{>} \left(\frac{2c}{\text{zMIC}}\right)^{\kappa} \tag{S4c}$$

For $\kappa \leq 1$, this inequality is never fulfilled since $\psi_{\min} < 0$, i.e. it always holds that $\mu(2c) \leq 2\mu(c)$. For $\kappa > 1$, it can be solved to yield:

$$\mu(2c) > 2\mu(c) \quad \Leftrightarrow \quad c < \left(\left(\frac{1}{2^{\kappa-1}} - 1\right) \cdot \frac{\psi_{\min}}{\psi_{\max}}\right)^{\frac{1}{\kappa}} \cdot \text{zMIC}. \tag{S5}$$

This indicates that a larger number of overlaps from same-drug doses could bring a benefit if $\kappa_A = \kappa_B > 1$ and drug concentrations are low. The situation during treatment is, of course, more complicated since antibiotic concentrations vary over time, and the cell population may contain resistant types. Figure S7C-F therefore show for all four cell types when presence of the two different drugs at concentrations $c_1$ and $c_2$ leads to a lower kill rate than presence of just one drug at concentration $c_1 + c_2$ (purple area in the figure), assuming $\kappa_A = \kappa_B = 2$. This again shows that for low antibiotic concentrations, overlaps of doses from the same drug lead to higher kill rates than overlaps of doses from different drugs for all types – even the double resistant type – except for the single resistant type that is resistant to that drug. This type is then managed by the administrations of the other drug.

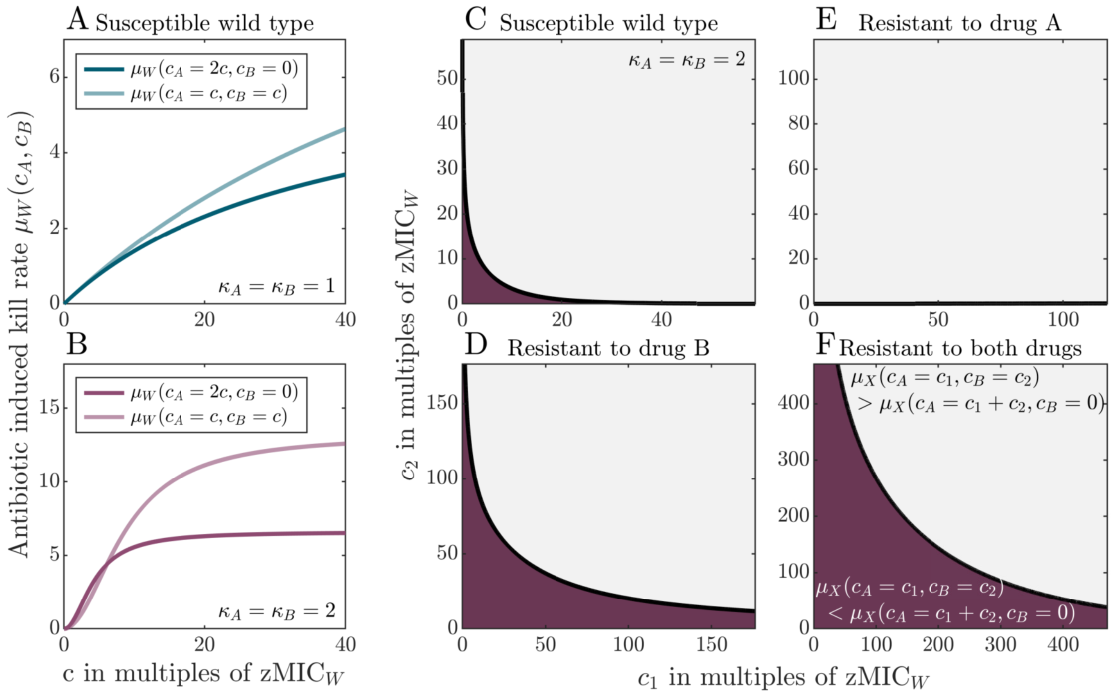


Figure S7: Comparison of the antibiotic-induced kill rates for the overlap of doses from the same drug and doses from different drugs. Panels A and B show the pharmacodynamic functions (Eq. (2)) of the susceptible wild type if either one drug is present at concentration $2c$ (and the other one absent) or if both drugs are present at concentration $c$ each. Panel A considers a drug pair with $\kappa_A = \kappa_B = 1$ and Panel B a drug pair with $\kappa_A = \kappa_B = 2$. Panels C-F show for all cell types when combining two drugs at concentrations $c_1$ and $c_2$ leads to a higher kill rate than having one drug at concentration $c_1 + c_2$ (grey area), considering a drug pair with $\kappa_A = \kappa_B = 2$. For the regions marked in lilac, a high concentration of one antibiotic leads to a higher kill rate than combining different drugs at lower concentrations.

#### S2.4 The effect of collateral resistance on treatment efficiency

Resistance to one drug can lead to an increase in susceptibility to another drug (collateral sensitivity) but also to an increase in resistance, which is called collateral resistance. Barbosa et al. [15] found that among experimental replicates, some strains that evolved resistance to a given drug displayed collateral resistance and other strains collateral sensitivity to another drug. In this section, we briefly consider the effect of collateral resistance on the treatment efficiency of sequential regimens. Figure S8 shows that unlike collateral sensitivity, collateral resistance has an effect even for rapid cycling regimens.

Similar to collateral sensitivity, these observations can be explained by the single resistant sub-population size in the treatments with different cycling frequencies. For slow regimens, the sub-population size is larger than for fast regimens. Hence a slight increase in resistance can make a visible difference in treatment efficiency - the same hold for the drug concentrations. The single resistant types can grow better at lower drug concentrations, resulting in a larger subpopulation than at high drug concentrations. Unlike collateral sensitivity, collateral resistance has a visible effect on fast cycling regimens, as it increases the first effective concentrations. The comparison shows that, even for fast cycling regimens, higher drug concentrations are needed for treatment success when resistance to one drug confers resistance to the other.

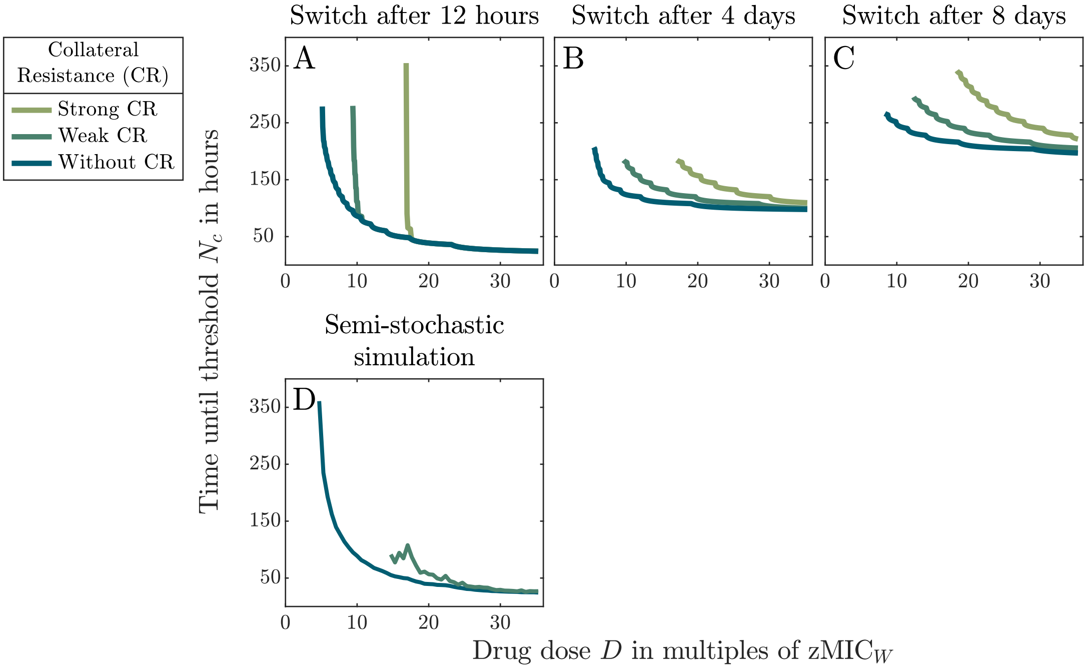


Figure S8: Time until the threshold $N_c$ is reached by all sub-populations for three different treatments with and without collateral resistance for a drug pair with $\kappa_A = \kappa_B = 1$. Panels A-C show results from the deterministic ODE model (1). Drugs get switched every 12 hours (**Panel A**), every four days (**Panel B**), or every eight days (**C**) . **Panel D** shows the results of semi-stochastic simulations for the treatment in which the drugs are switched every 12 hours. With weak collateral resistance, resistance to one drug increases resistance to the other drug (the respective zMIC value) by a factor of two, and with strong collateral resistance by a factor of four.

#### S2.5 Collateral sensitivity in the lab

Figure S9 shows the treatment efficiency in the lab when single resistant types display strong collateral sensitivity to the other drug. We see that the most rapid cycling remains optimal. Figure S10 further displays the direct comparison of the treatment efficiencies between drugs for which single resistant types display strong collateral sensitivity and drugs for which they do not at three different cycling frequencies.

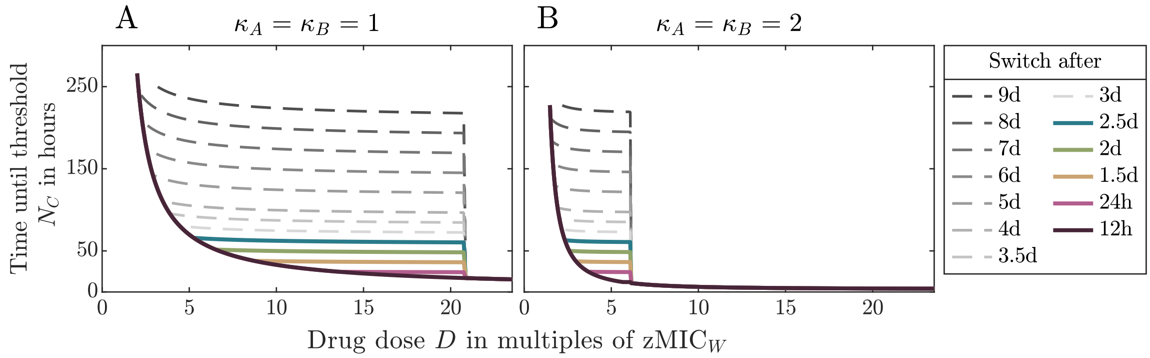


Figure S9: Treatment efficiency of different cycling regimens with drugs displaying collateral sensitivity in the lab. Panel A shows for $\kappa_A = \kappa_B = 1$ and Panel B for $\kappa_A = \kappa_B = 2$ the time until the threshold $N_c$ is reached by all sub-populations for different treatments across a wide range of concentration. In each treatment, strong collateral sensitivity is present.

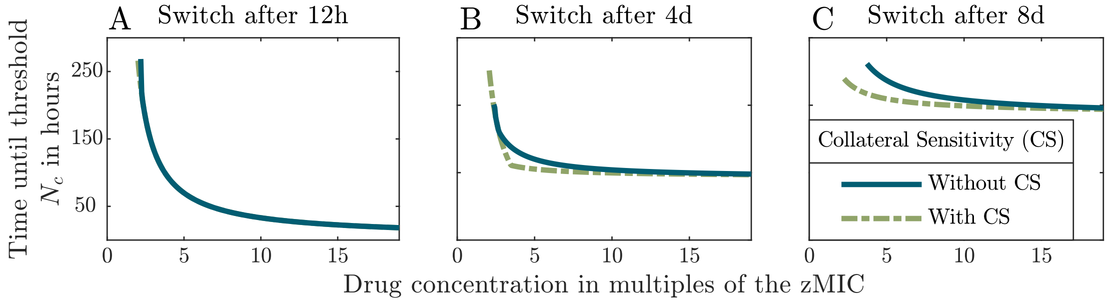


Figure S10: Treatment efficiency of drugs with and without strong collateral sensitivity in the lab for three different cycling frequencies. The graphs display the time until the threshold $N_c$ is reached by all sub-populations for drugs with $\kappa_A = \kappa_B = 1$.

##### S2.6 Treatment efficiency with and without collateral sensitivity and drug-drug interactions

In this section, we provide additional data for the treatment efficiency under various settings.

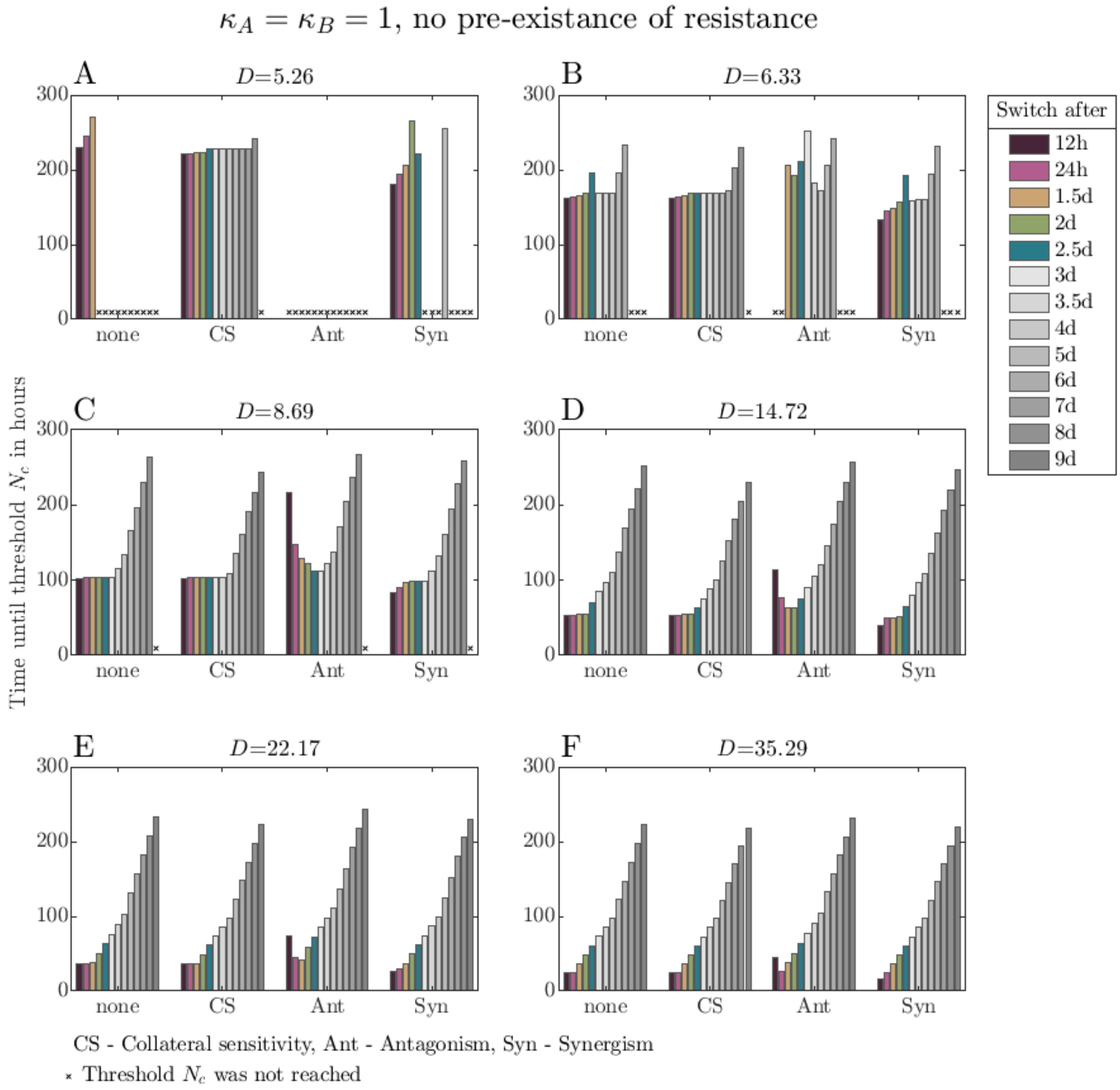


Figure S11: Comparison of treatment efficiencies for specific drug pairs with $\kappa_A = \kappa_B = 1$ with and without collateral sensitivity and drug-drug interactions. Each panel compares for some dose $D$ the time until all bacterial sub-population sizes have dropped below the threshold $N_c$ for different cycling frequencies under either Bliss independence without collateral sensitivity (none), under Bliss independence with collateral sensitivity (CS), under synergistic drug-drug interactions (Syn) or under antagonistic drug-drug interactions (Ant). The figure shows the inferiority of very rapid cycling over slower cycling under antagonistic interactions and an increase in the treatment efficiency of rapid cycling regimens under synergistic interactions. Collateral sensitivity improves the treatment efficiency of slow regimens. For low drug concentrations (Panels A and B), the dependency of the treatment efficiency on the cycling frequency looks erratic. This is a consequence of the deterministic nature of the model, similar to the effects described in SI section ??.

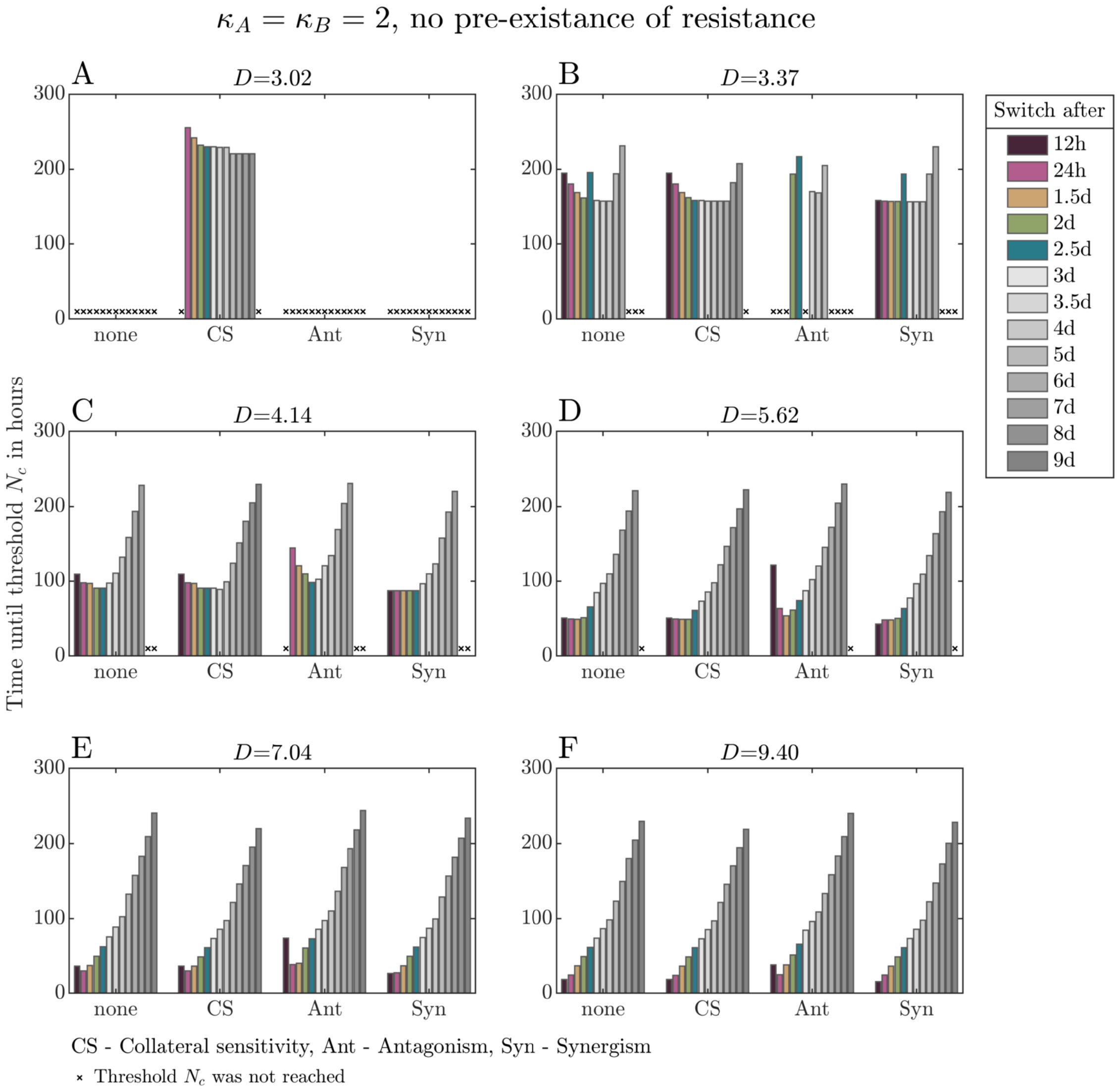


Figure S12: Comparison of treatment efficiencies for specific drug pairs with $\kappa_A = \kappa_B = 2$ with and without collateral sensitivity and drug-drug interactions. Each panel compares for some dose $D$ the time until all bacterial sub-population sizes have dropped below the threshold $N_c$ for different cycling frequencies under either Bliss independence without collateral sensitivity (none), under Bliss independence with collateral sensitivity (CS), under synergistic drug-drug interactions (Syn) or under antagonistic drug-drug interactions (Ant). The figure shows the inferiority of very rapid cycling over slower cycling under antagonistic interactions and an increase in the treatment efficiency of rapid cycling regimens under synergistic interactions. Collateral sensitivity improves the treatment efficiency of slow regimens. For low drug concentrations (panels A and B), the dependency of the treatment efficiency on the cycling frequency looks erratic. This is a consequence of the deterministic nature of the model, similar to the effects described in SI section ??.

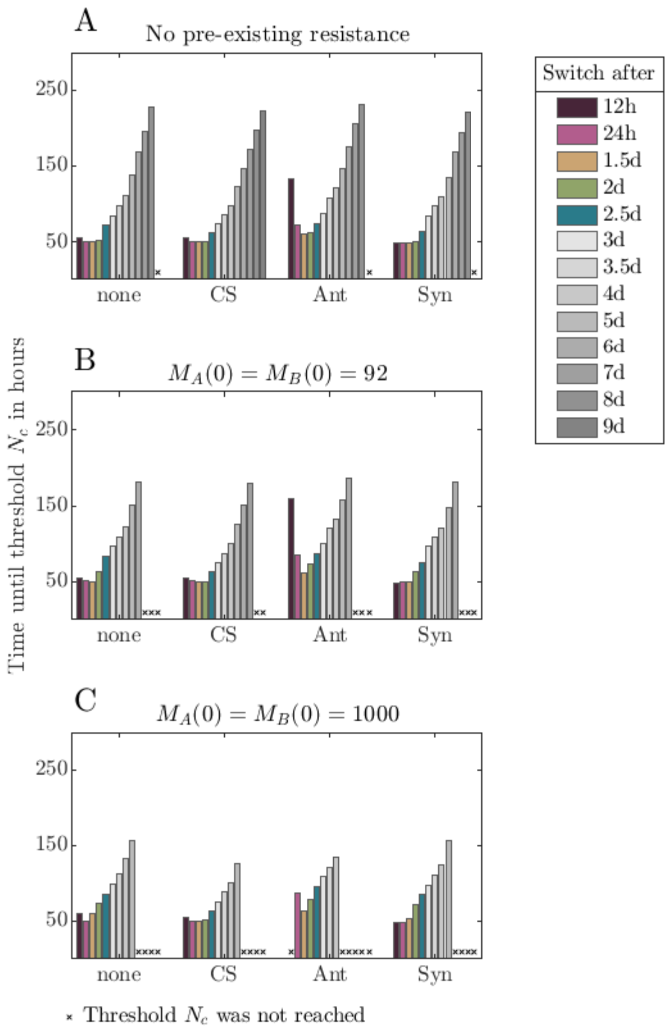


Figure S13: Comparison of treatment efficiencies for specific drug pairs with $\kappa_A = \kappa_B = 1$ with and without collateral sensitivity and drug-drug interactions and different initial populations. Each panel compares for dose $D = 5.43 \cdot \mathrm{zMIC}_W$ the time until the all sub-population sizes have dropped below the threshold $N_c$ for different cycling frequencies under either Bliss independence without collateral sensitivity (none), under Bliss independence with collateral sensitivity (CS), under synergistic drug-drug interactions (Syn) or under antagonistic drug-drug interactions (Ant). Panel A assumed no pre-existence of single resistance in the starting population at time $t = 0$, Panel B and C assume 90 and 1000 single resistant bacteria of each type.

#### S2.7 Effect of the Hill coefficient on selection of resistance for sub-inhibitory concentrations

While reviewing the literature regarding the effect of the Hill coefficient $\kappa$ on the treatment efficiency, we found two different conclusions. On the one hand, drugs with a large Hill coefficient were found to reduce the probability of resistance evolution and to lead to less treatment failure during mono-therapy [16]. On the other hand, drugs with low $\kappa$ were found to delay the fixation time of double mutants in sequential therapy [14].
As described in the main text, we find that treatments using drugs with high Hill coefficients are more efficient for the treatment of slowly replicating bacteria (note that the two studies cited above consider rapidly replicating bacteria, which we discuss further below in SI section S3). But we were only looking at concentrations that, anyway, result in a treatment success. Figure S14 compares the cell numbers of the double resistant type for a drug pair with $\kappa_A = \kappa_B = 1$ and a drug pair with $\kappa_A = \kappa_B = 2$ throughout the time course of a fast sequential treatment under concentrations not resulting in the clearance of the infection. The cell numbers are measured at the end of each administration period (one hour before the next dose is administered). Panels A and B show the cell numbers of the double resistant mutant for treatments with drug pairs with $\kappa_A = \kappa_B = 1$ and $\kappa_A = \kappa_B = 2$ respectively. The grey area shows regions in which the cell numbers remain below $N_c = 1$. Comparing these two panels with each other shows that the range of drug concentrations selecting for double resistance is much smaller for the drug pair with $\kappa_A = \kappa_B = 2$. This aligns with the observation that a Hill coefficient decreases the mutation selection concentration [17]. Panel C shows the difference between the cell numbers of the double resistant type for the two treatments and highlights regions in which use of either drug pair leads to a higher cell number (green: the drug pair with $\kappa_A = \kappa_B = 2$ leads to a higher number of double resistant cells, lilac: the drug pair with $\kappa_A = \kappa_B = 1$ leads to a higher number of double resistant cells) or regions in which the numbers are below $N_c = 1$ in both cases. The plot shows that there is indeed a range of concentrations in which treatment with the drug pair with $\kappa_A = \kappa_B = 2$ leads to higher cell numbers of the double resistant type. The effect of $\kappa_A = \kappa_B$ is thus concentration-dependent. We only show the results for the fast regimen, but the observations hold true for the other considered regimens as well.

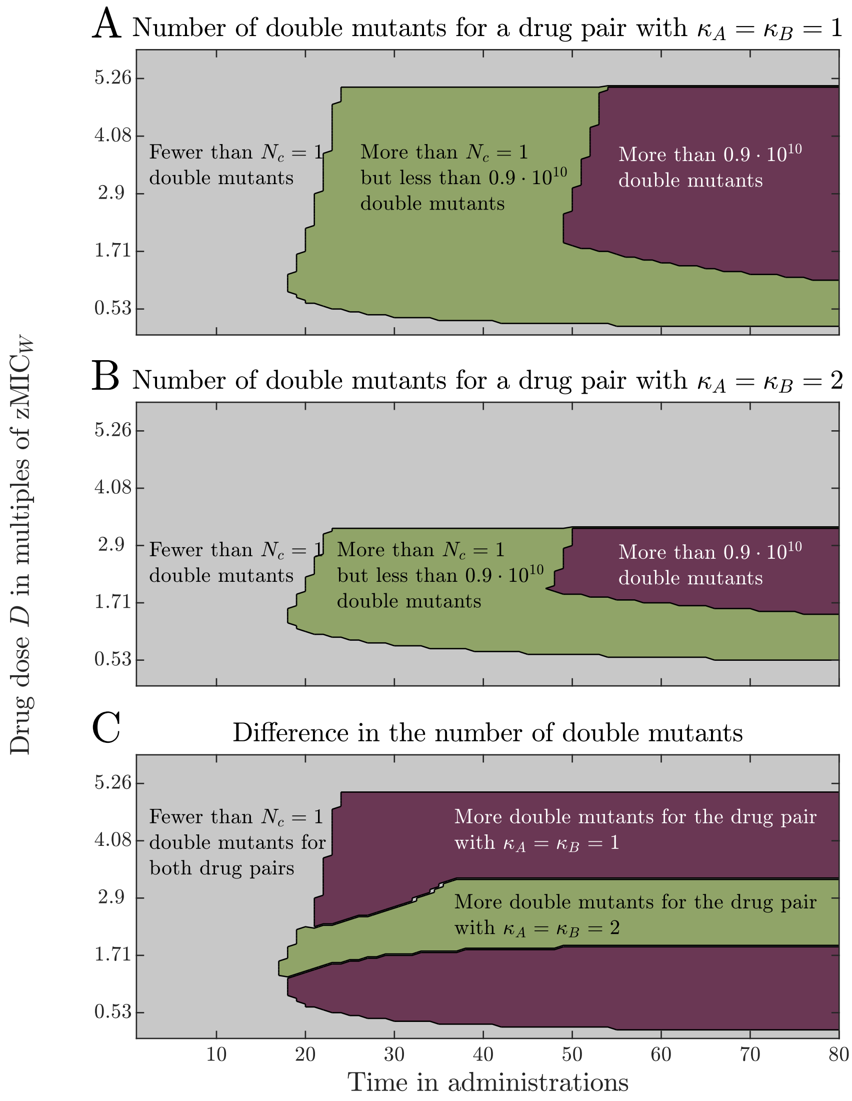


Figure S14: Effect of the Hill coefficients $\kappa_A = \kappa_B$ on the growth of the double resistant mutant for sub-inhibitory concentrations during a fast sequential regimen. The cell numbers are measured at the end of each administration period one hour before the next dose is given ($x$-axis) and for different drug concentrations ($y$-axis). Panels A and B show the cell numbers of the double resistant type for treatment with drug pairs with $\kappa_A = \kappa_B = 1$ and $\kappa_A = \kappa_B = 2$ respectively, divided in three categories: fewer than $N_c = 1$ cells (grey), cell numbers above $N_c$ but below $0.9 \cdot 10^{-9}$ (green), and cell numbers above $0.9 \cdot N_0$ (lilac). Panel C shows the difference in the double mutant cell numbers between treatments with the two drug pairs. The grey color indicates regions in which the cell numbers are below $N_c = 1$ for both treatments, the lilac color indicates regions in which the cell numbers are larger for a treatment with a drug pair with $\kappa_A = \kappa_B = 1$, and the green color indicates regions in which the cell numbers are larger for a treatment with a drug pair with $\kappa_A = \kappa_B = 2$.

#### S2.8 The effect of drug-drug interactions on selection of resistance for sub-inhibitory concentrations

Results for the effect of drug-drug interactions on combination therapy showed that synergism can lead to an increased clearance of the bacterial infection [15], but can also accelerate the evolution of resistance [18]. Similar to the effect of the Hill coefficient discussed in section S2.7, we can observe both effects in our model. As shown in the main text, synergism reduces the time to suppress all bacterial sub-populations below the threshold $N_c$. But here again, we only consider concentrations that are high enough to lead to a treatment success. In Figure S15, we explore the effect of drug-drug interactions on the growth of double resistant bacteria during a fast sequential regimen for sub-inhibitory concentrations. Each plot displays the difference between the cell numbers for two types of interaction (Bliss independence, synergism and antagonism), to identify regions in which either of the interactions leads to higher cell numbers of the resistant type. The comparison of synergism with Bliss independence (Panel A) shows that there is a concentration range in which synergism selects more strongly for double resistance. This can also be observed for the comparison between synergism and antagonism (Panel C). Bliss independence selects more strongly for double resistance than antagonism in some concentration ranges (Panel B). Similar to what was discussed by Torella et al. [19], we can explain these findings by the competition between the strains. For very low concentrations, which cannot even inhibit the growth of the susceptible type, the susceptible type will out-compete the other types. For high enough concentrations, the susceptible type is suppressed so quickly that it does not generate much resistance (for antagonism, much higher concentrations are needed for this). At intermediate concentrations, the resistant types profit from competitive release, but the extent depends on the drug-drug interaction and is greatest for synergism and lowest for antagonism.

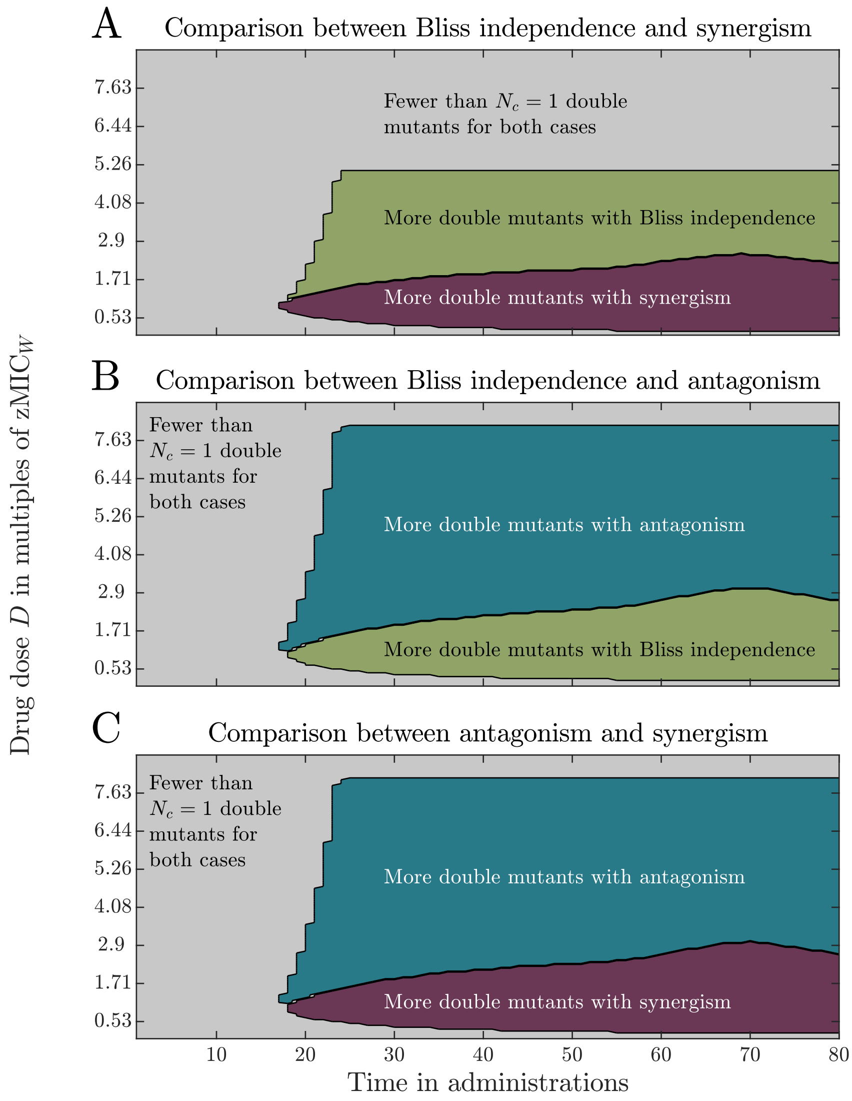


Figure S15: Effect of drug-drug interactions on the growth of double resistant mutants for sub-inhibitory concentrations during a fast sequential regimen. The cell numbers are measured at the end of each administration period one hour before the next dose is given ($x$-axis) and for different drug concentrations ($y$-axis). The panels display the differences in double mutant cell numbers for the comparison between synergism and Bliss independence (Panel A), antagonism and Bliss independence (Panel B), and synergism and antagonism (Panel C). Grey areas indicate areas in which the cell numbers are below $N_c = 1$ for both drug pairs, green color indicates regions in which Bliss independence leads to higher cell numbers, lilac color indicates regions in which synergism leads to higher cell numbers, and blue color indicates regions in which antagonisms leads to higher cell numbers.

### S3 Results for the treatment of rapidly replicating bacteria

#### S3.1 Treatment efficiency for rapidly replicating bacterial populations

In the main text, we assume that cells are replicating slowly (doubling time of $\ln(2)/\psi_{\max}^{W} \approx 7.88\ h$) to account for the fact that the body of the patient might not provide optimal growth conditions such as those supplied in laboratory experiments. In this section, we will discuss results for the treatment of rapidly replicating bacterial populations. We now assume a ten times higher growth rate than in the main text (doubling time of ca. 47.3 minutes).
We further compare the results with a second lab model that includes a population bottleneck after each growth period of one drug administration to model the dilution step of serial transfer experiments. The population is reduced to a hundredth of its size before the administration of the next drug. The bottleneck occurs in each implementation at the time point at which the drug is administered, hence the new reduced cell number is used once the next drug is present. In the deterministic model, this can be achieved by solving the ODE-system numerically for every administration separately and using each time the reduced population as starting population (except for the very first drug administration). For the semi-stochastic model, the bottleneck is implemented as follows: as the time steps are not equidistant, the time point of administration might not be reached exactly. To be able to reduce the population size correctly, we need to know how large the population at the time point of drug administration would be, before the reduction. We therefore check in every iteration whether by adding the new time step, the time point of administration would be exceeded. If by adding the time step $\tau$ the time point of administration is crossed, we know that the stochastic sub-populations will not change in cell number until the time of administrations (i.e. the next event will only occur after the time point of drug administration). We therefore only need to update the deterministic sub-populations. We then sample randomly from all sub-populations and set the new time point and new cell numbers at the time point of administration.

#### S3.2 Differences in growth rates (pharmacodynamics) between rapidly and slowly replicating populations

Before going into the details of the treatment comparison for rapidly replicating bacteria, we will give an overview on how the rate of the replication influences the concentration-dependent bacterial growth rates (the pharmacodynamics). In the absence of antibiotics, rapidly replicating bacteria have a higher growth rate than slowly replicating bacteria. This holds for all concentrations below the zMIC$_W$ (see Figure S16). When the concentration exceeds the zMIC$_W$, the growth rate of rapidly replicating wild-type cells is lower than that of slowly replicating wild-type cells (Panel A), i.e. rapidly replicating wild-type cells are more efficiently killed at concentrations exceeding the zMIC$_W$ than slowly replication wild-type cells. For resistant mutants, the MIC is substantially higher (here $28{\cdot}\mathrm{zMIC}_W$), and the curves are shifted to the right (Panel B). This means that at concentrations above the zMIC$_W$ but below the MIC of the resistant types, rapidly replicating wild-type cells grow worse than slowly replicating wild-type cells, but rapidly replicating resistant types have a higher growth rate than slowly replicating resistant types. Above the mutant MIC, rapidly replicating resistant types are again more efficiently killed than slowly replicating resistant types by the drug the mutants are resistant to. These observations are important for the results in the following sections.

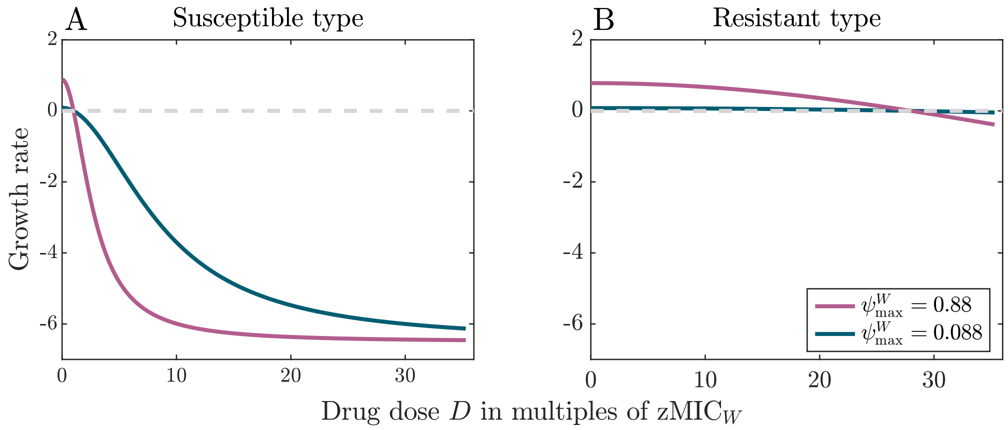


Figure S16: Pharmacodynamic functions for slowly and rapidly replicating bacteria. Panel A compares the growth rates of slowly replicating and rapidly replicating wild-type cells and Panel B the growth rates of single resistant bacteria. The parameter values used here are according to Table 1.

##### S3.2.1 Rapid cycling is always optimal for rapidly replicating populations

For rapidly replicating bacteria, as shown in Figure S17, the most rapid cycling is effective at lower drug doses than any slower cycling regimen and leads to the fastest treatment success (see Figures S18 and S19 for results from semi-stochastic simulations). This can be observed in every model (patient, lab without and lab with bottleneck) and for both $\kappa_A = \kappa_B = 1$ and $\kappa_A = \kappa_B = 2$.

Although rapidly replicating wild-type bacteria are eradicated much more efficiently than slowly replicating ones (see example dynamics in Figure S20), higher drug concentrations are needed to control the single-resistant sub-populations and to prevent the occurrence of double resistance if the intrinsic replication is high than if it is low. Due to their rapid replication, the single resistant sub-populations reach high population sizes, which also increases the chance of double resistance (see Panel B). Once double resistance arises, it takes over the population. Only if the antibiotic concentration is high, the single resistant sub-populations can become eradicated before double resistance appears (see Panel D). These dynamics become even more pronounced for slower cycling regimens, as shown in Panel E-H, where treatment of rapidly replicating bacteria requires concentrations above the MIC of the single resistant types.

For slowly replicating bacteria, we find that at low antibiotic concentrations, slightly slower cycling is advantageous for patient treatment if $\kappa_A = \kappa_B = 2$ (see main text). We do not find this for rapidly replicating bacteria. The concentration for which same-drug overlaps lead to higher kill rates for the wild-type than overlaps of different drugs (see Eq. S5) are lower than the first effective concentration.

We further find for slowly replicating bacteria that the first effective concentration is lower for drug pairs with $\kappa_A = \kappa_B = 2$ than for those with $\kappa_A = \kappa_B = 1$. For rapidly replicating bacteria, the analysis of the deterministic ODE system predicts that this holds for the most rapid cycling and for very slow cycling regimens; for intermediate cycling regimens, drug pairs with $\kappa_A = \kappa_B = 1$ require lower concentrations for treatment success. Example dynamics for such an intermeditae regime (switch every 1.5 days) are shown in Figure S21. Although the wild-type population gets eradicated faster for drugs with $\kappa_A = \kappa_B = 2$ than for drugs with

$\kappa_A = \kappa_B = 1$ (see also Figure 1E), the single resistant sub-populations grow better for drugs with a larger Hill coefficient when treating below the MIC of resistant types (see Figure S22). This leads to the occurrences of double resistance for rapidly replicating population treated with drugs having a large Hill coefficient (see Figure S21 D).
Nevertheless, also for rapidly replicating bacteria, the time until all sub-population sizes are suppressed below $N_c$ is lower for drug pairs with $\kappa_A = \kappa_B = 2$ than for $\kappa_A = \kappa_B = 1$ once the concentration is high enough to achieve treatment success, which can be deduced from the lower concentration at which the sudden drop occurs in the deterministic model (Figure S17) and can also be directly seen in the results from the semi-stochastic simulations (Figure S18).

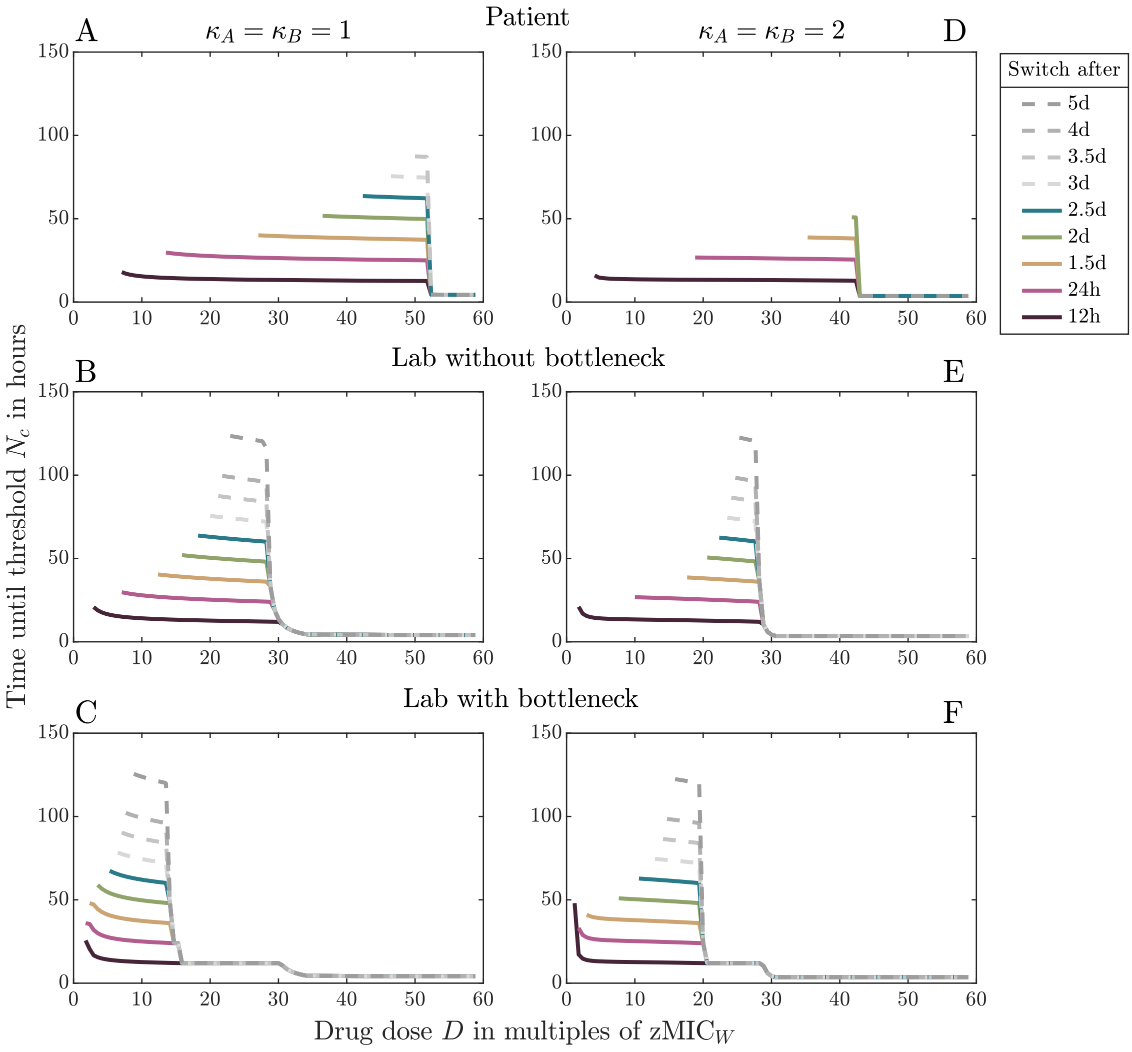


Figure S17: Treatment efficiency of treatments of rapidly replicating bacteria for drugs with $\kappa_A = \kappa_B = 1$ (Panel A-C) and $\kappa_A = \kappa_B = 2$ (Panel D-F). The first row (Panels A and D) show the treatment efficiency for the patient model and the second and third rows for the lab model without (Panels B and E) and with population bottlenecks (Panels C and F).

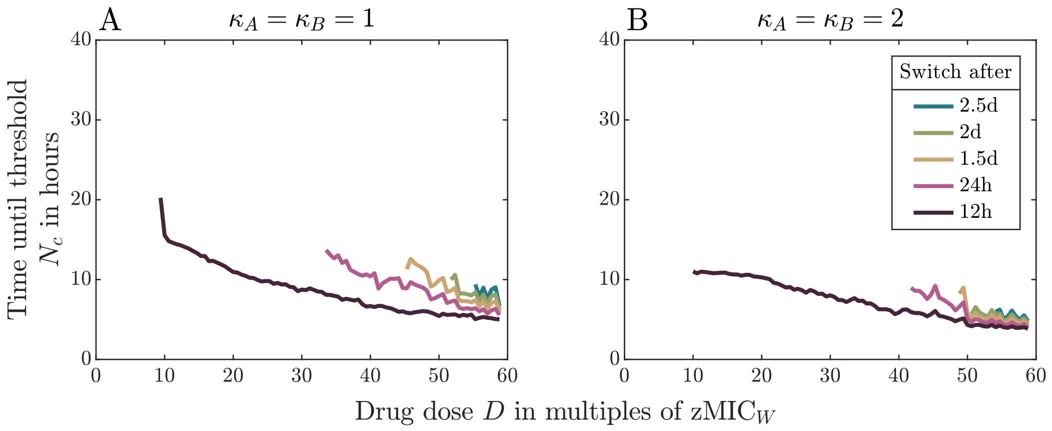


Figure S18: Results from semi-stochastic simulations for the patient model for treatments of rapidly replicating bacteria.

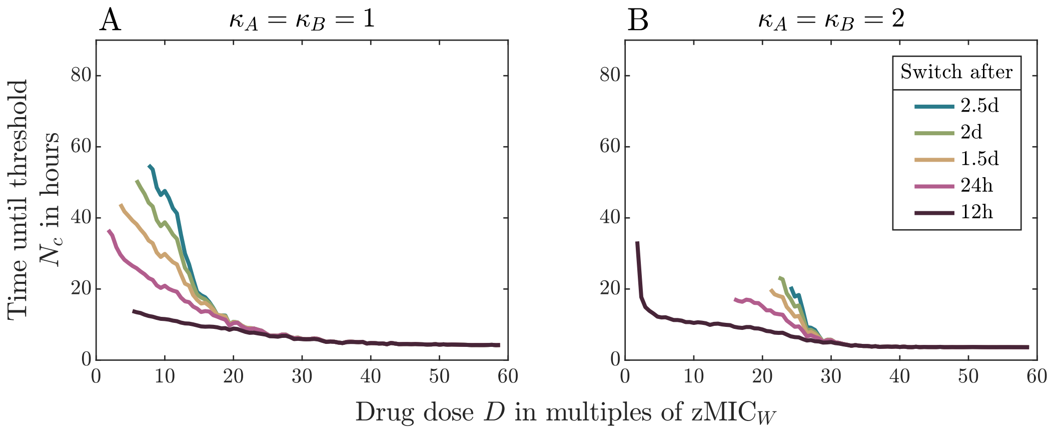


Figure S19: Results from semi-stochastic simulations for the lab model with population bottlenecks for treatments of rapidly replicating bacteria.

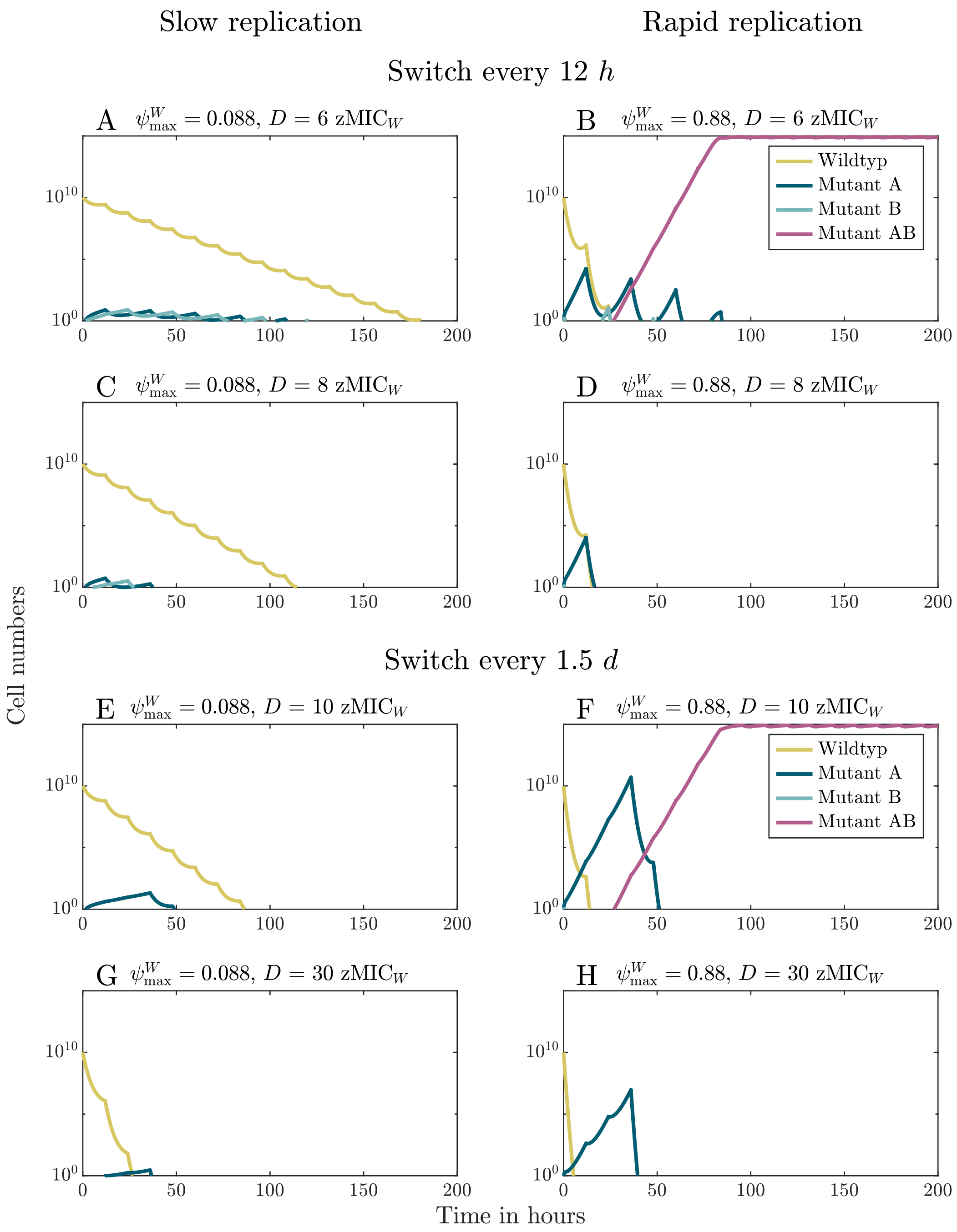


Figure S20: Example dynamics for the treatment of slowly and rapidly replicating bacteria for two different cycling regimens. The drugs have Hill coefficients $\kappa_A = \kappa_B = 1$.

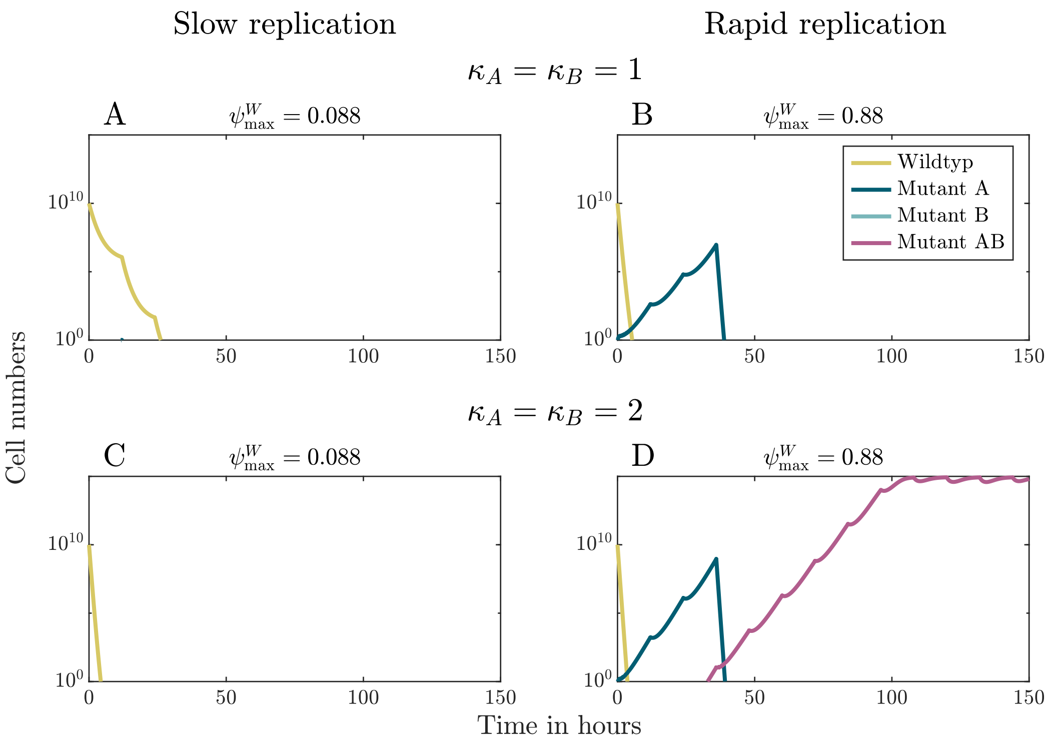


Figure S21: Example dynamics for the treatment of slowly and rapidly replicating bacteria at a specific drug concentration for drugs with different Hill coefficients. The drug is switched every 1.5 days. The drugs are applied at dose $D = 7 \cdot \text{zMIC}_W$.

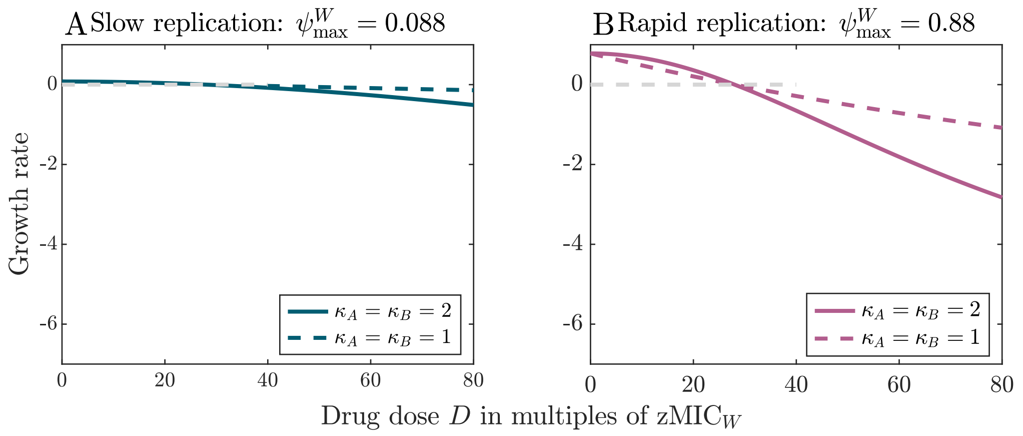


Figure S22: Pharmacodynamic functions for slowly and rapidly replicating resistant bacteria for drugs with different Hill coefficients. The parameter values used here are according to Table 1.

##### S3.2.2 Collateral effects and drug-drug-interactions both have an effect on rapid cycling regimens

Overall, we find that effects of collateral sensitivity on the treatment efficiency are very small if bacteria divide rapidly (see Figures S23, S24A, S25).
Figure S23A-C show that for rapidly replicating bacteria unlike for slowly replicating ones, collateral sensitivity slightly increases the treatment efficiency of rapid cycling regimens at low concentrations (see Figures S24A and S25 for results of semi-stochastic simulations for the patient and the lab in the presence of periodic bottlenecks). Example dynamics at low concentrations are shown in Figure S26. For slowly replicating bacteria and rapid cycling, the time to suppress all sub-population sizes below the threshold $N_c$ depends on the wild-type sub-population and is thus not influenced by collateral sensitivity effects (Panels A and C). For rapidly replicating bacteria, in contrast, resistant bacteria may become prevalent and influence the time to reach $N_c$. Especially, collateral sensitivity can then prevent the occurrence of double resistance (compare Panels B and D).
For slow cycling regimens (Figure S23D-F), collateral sensitivity only increases the treatment efficiency in the lab model with bottlenecks, but not in the other two models. The effect for the lab model with bottlenecks is not visible in results from semi-stochastic simulations within the considered range of concentrations (Figure S25). While (unlike for slowly replicating bacteria) the sub-populations of single resistant bacteria can become large even at high concentrations (Figure S20 E-H), the concentrations required for treatment success in the patient model and the lab model without bottlenecks are so high that collateral sensitivity does not make a big difference in the growth rate itself and therefore cannot strongly improve the treatment efficiency (see Figure S27).
Drug-drug interactions can change the treatment efficiency, but the effect is less strong than for slowly replicating bacteria (Figure S28A and B; see Figure S29 for the pharmacodynamic functions). Since the effect of drug-drug-interactions is small compared to the differences in efficiency between the cycling regimens for fast replicating bacteria, antagonistic drug-drug-interactions do not shift the optimal cycling frequency to slower cycling. I.e., for antagonistic interactions, fast cycling is the optimal choice as shown in Figure S30.

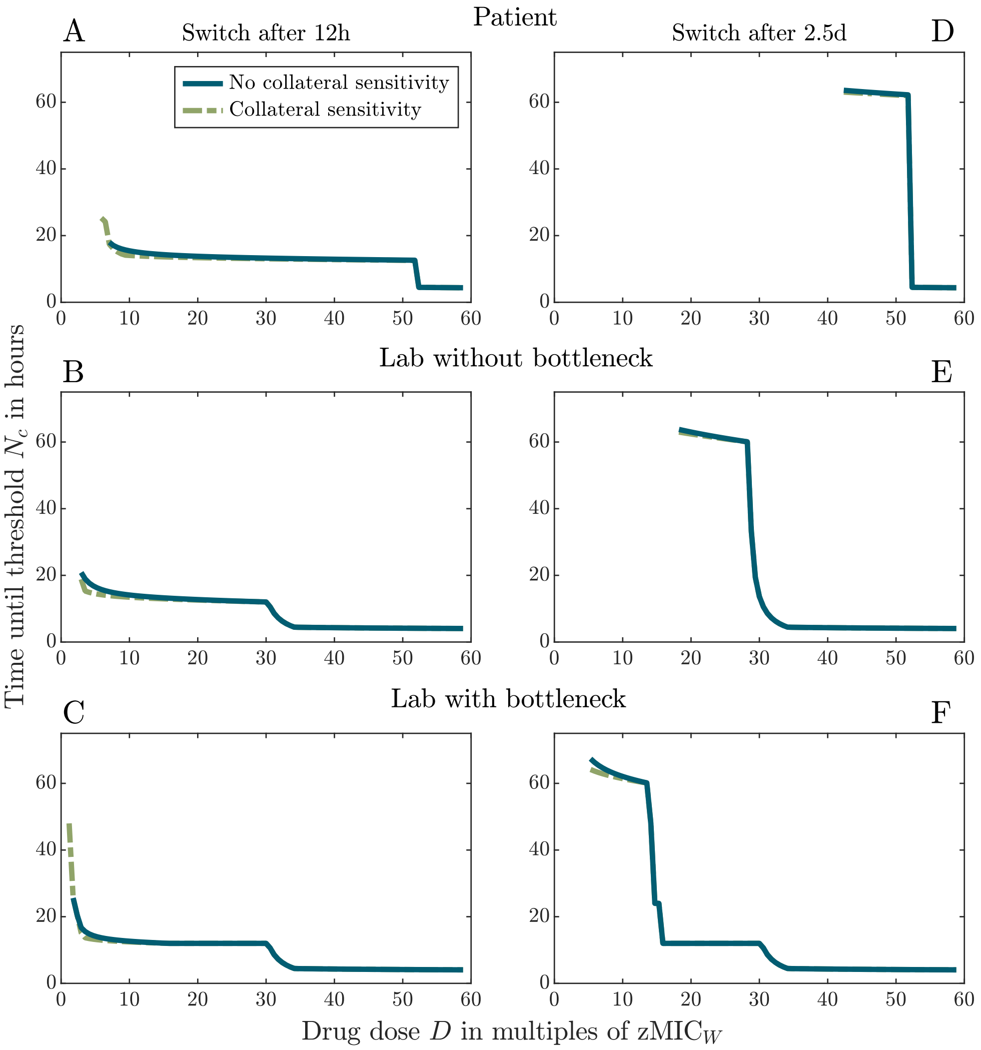


Figure S23: Effect of collateral sensitivity on the treatment of rapidly replicating bacteria for two cycling regimens.

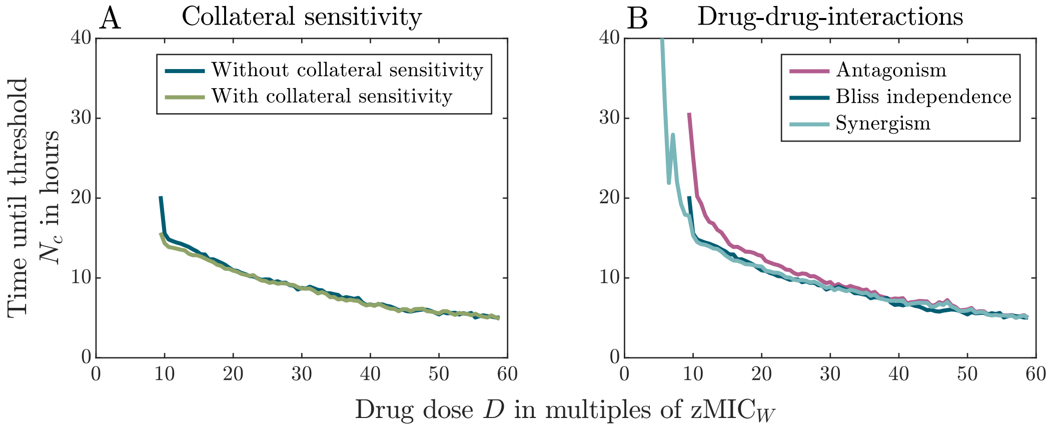


Figure S24: Results from semi-stochastic simulations for treatment of rapidly replicating bacteria in the patient with drugs displaying collateral sensitivity (Panel A) and drug-drug-interactions (Panel B). Drugs get switched every 12 hours (most rapid cycling).

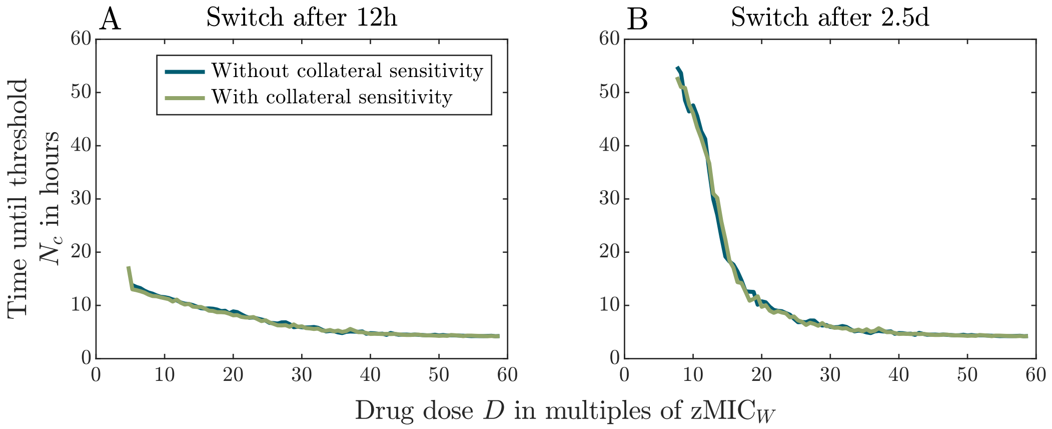


Figure S25: Results from semi-stochastic simulations for treatments of rapidly replicating bacteria in the lab model with population bottlenecks for drugs with and without collateral sensitivity. Panel A and Panel B display treatment in which the drugs are switched every 12 hours and every 2.5 days, respectively.

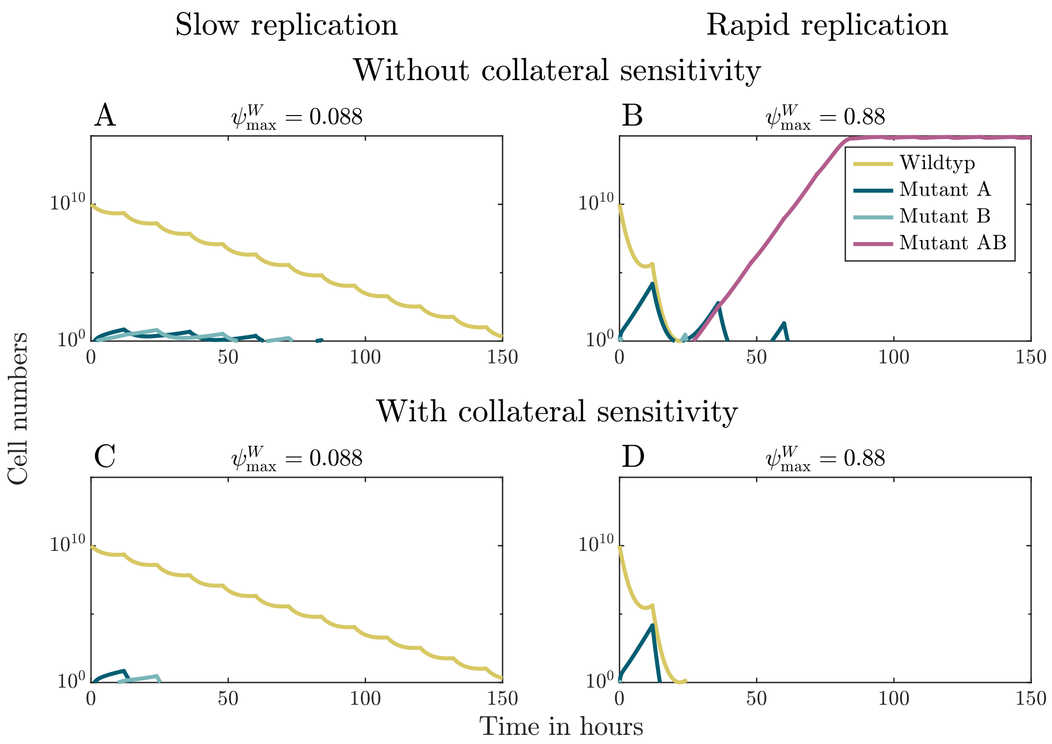


Figure S26: Example dynamics for the treatment of slowly and rapidly replicating bacteria using drugs with collateral sensitivity (Panels A and B) and without collateral sensitivity (Panels C and D) for the most rapid cycling regimen. The drug concentration is $D = 6.5 \cdot \text{zMIC}_W$, and the Hill coefficients are $\kappa_A = \kappa_B = 1$.

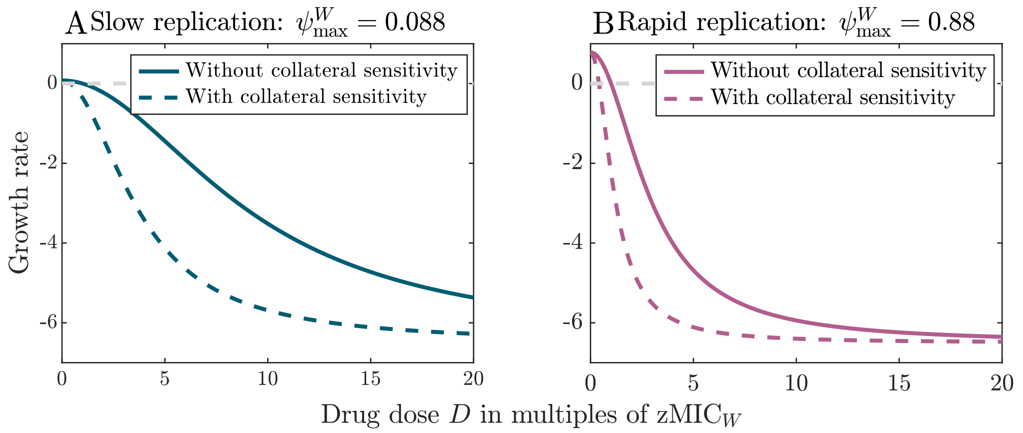


Figure S27: Pharmacodynamic functions for slowly and rapidly replicating bacteria in the presence and absence of collateral sensitivity. The parameter values used here are according to Table 1.

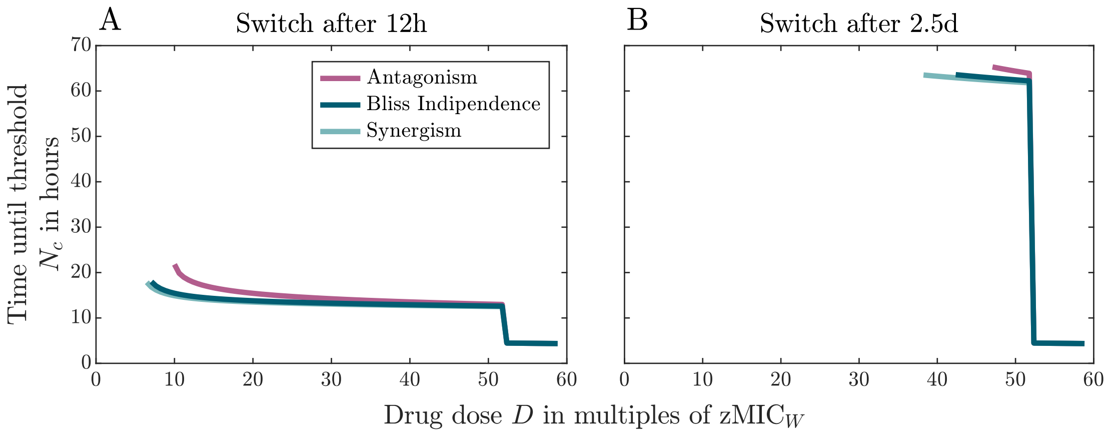


Figure S28: Effect of drug-drug-interactions on the treatment efficiency for rapidly replicating bacterial populations.

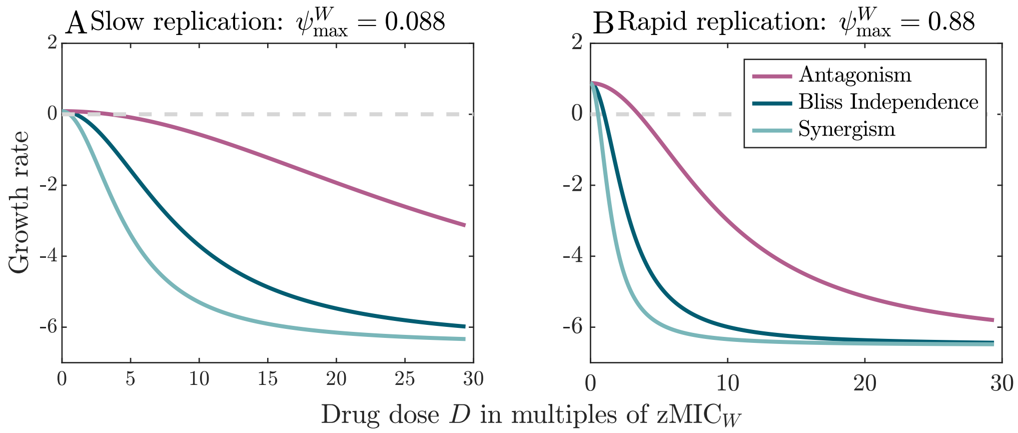


Figure S29: Pharmacodynamic functions for slowly and rapidly replicating bacteria with and without drug-drug-interactions. The parameter values used here are according to Table 1.

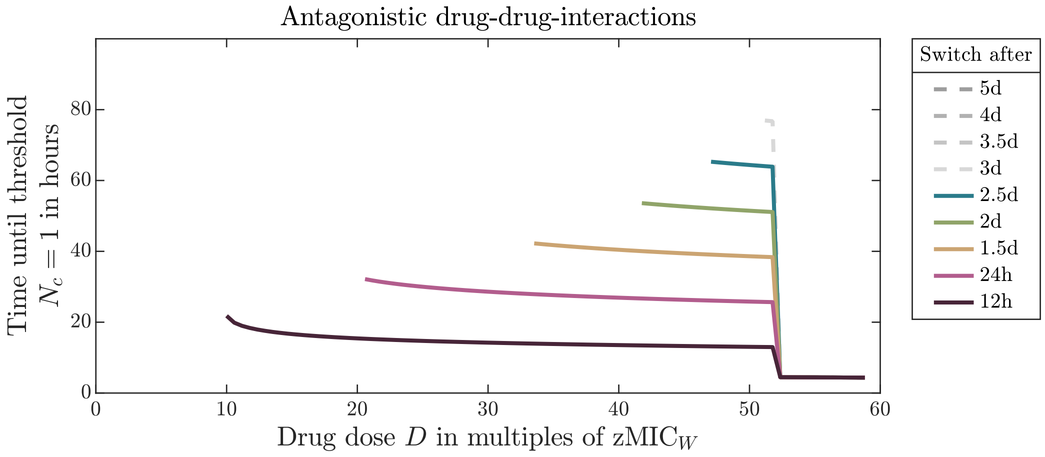


Figure S30: Effect of antagonistic drug-drug-interactions on the treatment efficiency for rapidly replicating bacteria in the patient for a range of cycling regimens.

#### References


(1) Wicha, S. G., Chen, C., Clewe, O., and Simonsson, U. S. H. (2017). A general pharmacodynamic interaction model identifies perpetrators and victims in drug interactions. *Nature Communications 8*, 1–11.

(2) Regoes, R. R., Wiuff, C., Zappala, R. M., Garner, K. N., Baquero, F., and Levin, B. R. (2004). Pharmacodynamic functions: a multiparameter approach to the design of antibiotic treatment regimens. *Antimicrobial Agents and Chemotherapy 48*, 3670–3676.

(3) Yeh, P. J., Hegreness, M. J., Presser Aiden, A., and Kishony, R. (2009). Drug interactions and the evolution of antibiotic resistance. *Nature Reviews Microbiology 7*, 460–466.

(4) Ankomah, P., and Levin, B. R. (2012). Two-drug antimicrobial chemotherapy: a mathematical model and experiments with *Mycobacterium marinum*. *PLoS Pathogens 8*, e1002487.

(5) Bollenbach, T. (2015). Antimicrobial interactions: mechanisms and implications for drug discovery and resistance evolution. *Current Opinion in Microbiology 27*, 1–9.

(6) Baym, M., Stone, L. K., and Kishony, R. (2016). Multidrug evolutionary strategies to reverse antibiotic resistance. *Science 351*.

(7) Singh, N., and Yeh, P. (2017). Suppressive drug combinations and their potential to combat antibiotic resistance. *The Journal of Antibiotics 70*, 1033–1042.

(8) Zimmer, A., Katzir, I., Dekel, E., Mayo, A. E., and Alon, U. (2016). Prediction of multidimensional drug dose responses based on measurements of drug pairs. *Proceedings of the National Academy of Sciences 113*, 10442–10447.

(9) Katzir, I., Cokol, M., Aldridge, B. B., and Alon, U. (2019). Prediction of ultra-high-order antibiotic combinations based on pairwise interactions. *PLoS Computational Biology 15*, e1006774.

(10) Lewis, P. A. W., and Shedler, G. S. (1979). Simulation of nonhomogeneous Poisson processes by thinning. *Naval Research Logistics Quarterly 26*, 403–413.

(11) Gillespie, D. T. (1976). A general method for numerically simulating the stochastic time evolution of coupled chemical reactions. *Journal of Computational Physics 22*, 403–434.

(12) Gillespie, D. T. (1977). Exact stochastic simulation of coupled chemical reactions. *The Journal of Physical Chemistry 81*, 2340–2361.

(13) Kiehl, T. R., Mattheyses, R. M., and Simmons, M. K. (2004). Hybrid simulation of cellular behavior. *Bioinformatics 20*, 316–322.

(14) Udekwu, K. I., and Weiss, H. (2018). Pharmacodynamic considerations of collateral sensitivity in design of antibiotic treatment regimen. *Drug Design, Development and Therapy 12*, 2249.

(15) Barbosa, C., Beardmore, R. E., Schulenburg, H., and Jansen, G. (2018). Antibiotic combination efficacy (ACE) networks for a *Pseudomonas aeruginosa* model. *PLoS Biology 16*, e2004356.

(16) Yu, G., Baeder, D. Y., Regoes, R. R., and Rolff, J. (2018). Predicting drug resistance evolution: insights from antimicrobial peptides and antibiotics. *Proceedings of the Royal Society B: Biological Sciences 285*, 20172687.

(17) Greenfield, B. K., Shaked, S., Marrs, C. F., Nelson, P., Raxter, I., Xi, C., McKone, T. E., and Jolliet, O. (2018). Modeling the emergence of antibiotic resistance in the environment: an analytical solution for the minimum selection concentration. *Antimicrobial Agents and Chemotherapy 62*, e01686–17.

(18) Hegreness, M., Shoresh, N., Damian, D., Hartl, D., and Kishony, R. (2008). Accelerated evolution of resistance in multidrug environments. *Proceedings of the National Academy of Sciences 105*, 13977–13981.

(19) Torella, J. P., Chait, R., and Kishony, R. (2010). Optimal drug synergy in antimicrobial treatments. *PLoS Computational Biology 6*, e1000796.
